# Liver lipid droplet cholesterol content is a key determinant of metabolic dysfunction-associated steatohepatitis

**DOI:** 10.1101/2025.02.25.640203

**Authors:** Ikki Sakuma, Rafael C. Gaspar, Ali R. Nasiri, Sylvie Dufour, Mario Kahn, Jie Zheng, Traci E. LaMoia, Mateus T. Guerra, Yuki Taki, Yusuke Kawashima, Dean Yimlamai, Mark Perelis, Daniel F. Vatner, Kitt Falk Petersen, Maximilian Huttasch, Birgit Knebel, Sabine Kahl, Michael Roden, Varman T. Samuel, Tomoaki Tanaka, Gerald I. Shulman

## Abstract

Metabolic dysfunction-associated steatohepatitis (MASH) represents a progressive form of steatotic liver disease which increases the risk for fibrosis and advanced liver disease. The accumulation of discrete species of bioactive lipids has been postulated to activate signaling pathways that promote inflammation and fibrosis. However, the key pathogenic lipid species is a matter of debate. We explored candidates using various dietary, molecular, and genetic models. Mice fed a choline-deficient L-amino acid-defined high-fat diet (CDAHFD) developed steatohepatitis and manifested early markers of liver fibrosis associated with increased cholesterol content in liver lipid droplets within 5 days without any changes in total liver cholesterol content. Treating mice with antisense oligonucleotides (ASOs) against *Coenzyme A synthase (Cosay)* or treatment with bempedoic acid or atorvastatin decreased liver lipid droplet cholesterol content and prevented CDAHFD-induced MASH and the fibrotic response. All these salutary effects were abrogated with dietary cholesterol supplementation. Analysis of human liver samples demonstrated that cholesterol in liver lipid droplets was increased in humans with MASH and liver fibrosis and was higher in PNPLA3 I148M (variants rs738409) than in HSD17B13 variants (rs72613567). Together, these data identify cholesterol in liver lipid droplets as a critical mediator of MASH and demonstrate that COASY knockdown and bempedoic acid are novel therapeutic approaches to reduce liver lipid droplet cholesterol content and thereby prevent the development of MASH and liver fibrosis.

**Significance Statement:** Metabolic dysfunction-associated steatohepatitis (MASH) is a progressive liver disease linked to fibrosis. The role of specific lipid species in its pathogenesis remains debated. Using dietary, molecular, and genetic models, we found that mice on a choline-deficient, high-fat diet (CDAHFD) developed steatohepatitis and early fibrosis, marked by increased cholesterol in liver lipid droplets within five days. Targeting COASY with antisense oligonucleotides or treating with bempedoic acid or atorvastatin reduced lipid droplet cholesterol and prevented MASH. However, dietary cholesterol supplementation negated these effects. Human liver samples confirmed elevated lipid droplet cholesterol in MASH and fibrosis, especially in PNPLA3 I148M carriers. These findings highlight cholesterol reduction as a potential MASH therapy.

## Introduction

Metabolic dysfunction-associated steatosis liver disease (MASLD) is the predominant chronic liver disease in the world, affecting approximately 25% of the global population (1, 2). The prevalence of MASLD is on the rise, paralleling a rise in obesity and type 2 diabetes. Metabolic dysfunction-associated steatohepatitis (MASH) increases the risk for advanced liver diseases, including cirrhosis, hepatocellular carcinoma, and liver-related death (2). While the U.S. Food and Drug Administration has recently approved resmetirom, a liver-directed, β-selective thyroid hormone receptor agonist, as a therapy for MASH with significant fibrosis only 29% of the study participants responded to this treatment in a recent Phase III clinical trial (3), demonstrating the need for additional new targets and therapies for MASH.

One of the major unanswered questions is what triggers the conversion of “benign” fatty liver disease to metabolic dysfunction-associated steatohepatitis (MASH). Triglycerides, the predominant lipids within hepatocytes, are neutral lipids that can be exported in VLDL particles or oxidized to support hepatocellular metabolism. Triglycerides reside within the core of lipid droplets, surrounded by a phospholipid monolayer and a host of proteins (e.g. lipases, transacylases and cofactors) that closely balance the influx and efflux of triglycerides from the lipid droplets. Though triglycerides are an important source of cellular energy, they are considered inert in terms of modulating cellular functions. Instead, other lipid species, such as fatty acids, cholesterol, lysophosphatidic acid, lysophosphatidylcholine, and ceramides, have all been implicated in causing hepatic inflammation and fibrosis (4–7). The choline-deficient L-amino acid-defined high-fat diet (CDAHFD) model has been advanced as a mouse model that mirrors the pathological progression of MASLD (steatosis ➾ hepatitis ➾ fibrosis) (8, 9). CDAHFD model displays MASH pathology within weeks, enabling a more expedited pathological evaluation (10). We postulated that pathogenic lipid species accumulate early in specific subcellular compartments in this model, contributing to organelle dysfunction and cellular injury. Thus, we measured candidate lipid species in key subcellular compartments in the livers of CDAHFD-fed C57BL/6J mice. We began these studies with a 5-day time course experiment of CDAHFD and observed a marked increase in cholesterol specifically in the liver lipid droplets within the first few days of treatment, which was associated with hepatic inflammation and markers of liver fibrosis. This observation informed our central hypothesis that cholesterol in liver lipid droplets triggers liver inflammation and fibrosis. We tested this hypothesis using cholesterol-lowering treatments (bempedoic acid, atorvastatin), cholesterol supplementation, and an antisense oligonucleotide (ASO) to knockdown the expression of Coenzyme A synthase (COASY). COASY generates CoA for acetyl CoA synthesis, a key substrate for cholesterol synthesis. To translate these findings to humans we examined the role of cholesterol in liver lipid droplets in humans with and without MASH/liver fibrosis as well the effects of established gene variant (HSD17B13) that have been shown to be protective against the development of MASH and liver fibrosis.

## Results

### Choline-deficient L-amino acid-defined high-fat diet acutely induces hepatic lipid accumulation followed by inflammation and fibrosis

Male C57BL/6J mice were divided into regular chow (RC) fed or CDAHFD fed groups. CDAHFD was administered daily for a duration of 1 to 5 days (Figure 1A). Plasma concentrations of ALT and AST increased within the first few days whereas plasma total cholesterol and triglycerides decreased in CDAHFD-fed mice (Figure 1B). Hematoxylin, eosin (HE), and BODIPY staining demonstrated that liver lipid droplets increased in size in a time-dependent manner (Figure 1C). CD68 staining revealed crown-like structures, which are macrophages surrounding and engulfing dying or dead hepatocytes with large lipid droplets and indicative of MASH (11). These structures were observed from day 3 of the CDAHFD diet onwards (Figure 1C and 1D). Filipin, which stains free cholesterol but not cholesterol esters, demonstrated an accumulation of free cholesterol within the lipid droplets as early as day 2 of the CDAHFD feeding (Figure 1C). These findings confirmed the development of steatohepatitis within 5 days of CDAHFD.

**Figure 1.**
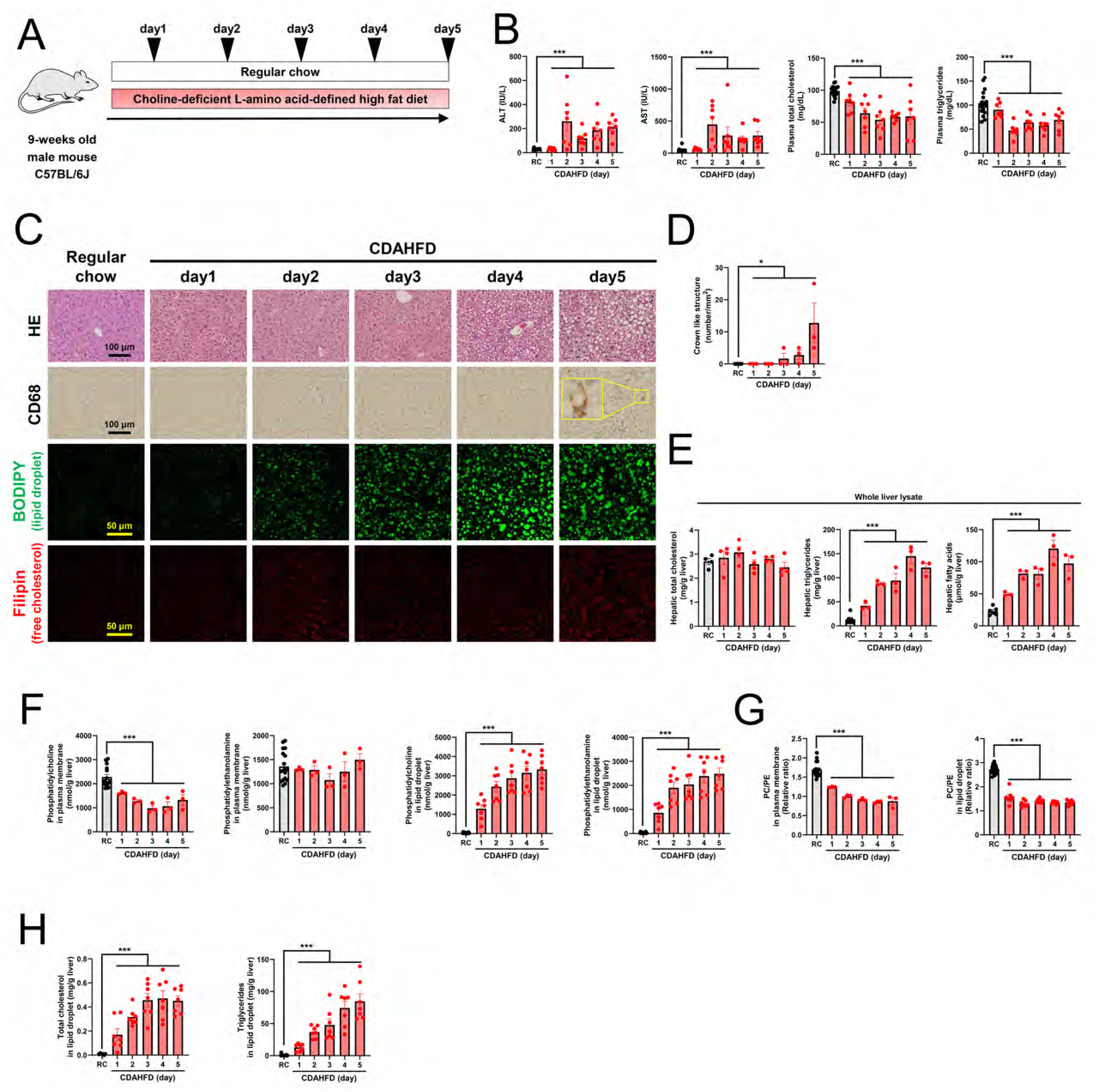
Choline-deficient L-amino acid-defined high-fat diet acutely induces hepatic lipid accumulation followed by inflammation and fibrosis. (A) Study design. C57BL/6J mice were divided into regular chow (RC) fed mice or choline-deficient, L-amino acid-defined, high-fat diet (CDAHFD) fed mice. CDAHFD was provided for 1, 2, 3, 4, and 5 days, respectively. (B) Plasma ALT, AST, total cholesterol and triglycerides levels. ALT and AST increased in CDAHFD-fed mice. Plasma total cholesterol and triglycerides decreased in CDAHFD-fed mice. (C) Liver sections were stained with hematoxylin, eosin (HE), CD68, BODIPY, and Filipin. CDAHFD induced liver lipid droplets time-dependently. CD68 staining revealed crown-like structures on day 5. BODIPY and Filipin staining demonstrated that free cholesterol was included in liver lipid droplets. (D) The number of crown-like structures increased in the liver of CDAHFD-fed mice time-dependently. (E) Total cholesterol, triglycerides, and fatty acids content in whole liver lysate. Total cholesterol did not change. Triglycerides and fatty acids increased in CDAHFD-fed mice time-dependently. (F) The levels of phospholipids in the liver. The level of phosphatidylcholine in the plasma membrane exhibited a decline starting on day 1. The level of phosphatidylethanolamine in the plasma membrane did not change. In liver lipid droplets, phosphatidylcholine and phosphatidyl ethanolamine levels increased from day 1. (G) The ratio of phosphatidylcholine (PC) to phosphatidylethanolamine (PE) in the plasma membrane and lipid droplet in the liver, respectively. PC/PE ratio in CDAHFD-fed mice was lower than RC-fed mice. (H) Total cholesterol and triglycerides in the liver lipid droplet increased in CDAHFD-fed mice from day 1. Data are presented as mean ± SEM. Groups were compared by Unpaired one-sided Student’s t-test.

We next quantified candidate lipid species implicated in the pathogenesis of MASH. Triglycerides and fatty acids in whole liver lysate increased in CDAHFD-fed mice in a time-dependent manner (Figure 1E). In contrast, total cholesterol content including free cholesterol and cholesterol esters in whole liver lysate did not change appreciably over the 5-day feeding period. Lower ratios of phosphatidylcholine (PC) to phosphatidylethanolamine (PE) can decrease membrane integrity and contribute to liver damage (12). To examine the relative concentrations of these two phospholipids, we isolated plasma membrane and lipid droplet fractions from whole liver tissues using ultracentrifugation, as described in previous studies and as depicted in Figure S1A (13, 14). In the plasma membrane, PC concentrations declined on day 1, reached a nadir on day 3 and showed an upward trend on days 4 and 5 (Figure 1F). The level of PE in the plasma membrane did not change (Figure 1F). In contrast, within the liver lipid droplets, both PC and PE concentrations increased starting from day 1 (Figure 1F). Despite the differences in PC and PE absolute concentrations in these discrete subcellular compartments, the ratio of PC to PE in CDAHFD-fed mice compared to regular chow-fed mice decreased in plasma membrane and lipid droplets (Figure 1G).

We next evaluated total cholesterol and triglycerides in liver lipid droplets. Total cholesterol and triglycerides in liver lipid droplets increased in CDAHFD-fed mice from day 1 (Figure 1H). LC-MS/MS analysis consistently verified that free cholesterol and cholesterol ester in liver lipid droplets increased (Figure S1B).

Taken together, CDAHFD feeding acutely induced hepatic lipid accumulation followed by inflammation and fibrosis. Total cholesterol in whole liver lysate did not track with inflammation and fibrosis markers, but total cholesterol in liver lipid droplets tracked with inflammation and fibrosis.

### Choline-deficient L-amino acid-defined high-fat diet induces hepatocytes death possibly via lysosomal membrane disruption

Next, we compared CDAHFD-fed mice with RC-fed mice after one week of feeding (Figure 2A). Consistent with prior findings CDAHFD-fed mice did not induce obesity or hyperglycemia (8) and there were no differences in percent body fat or lean body mass as assessed by ^1^H NMR in (Figure S2A).

**Figure 2.**
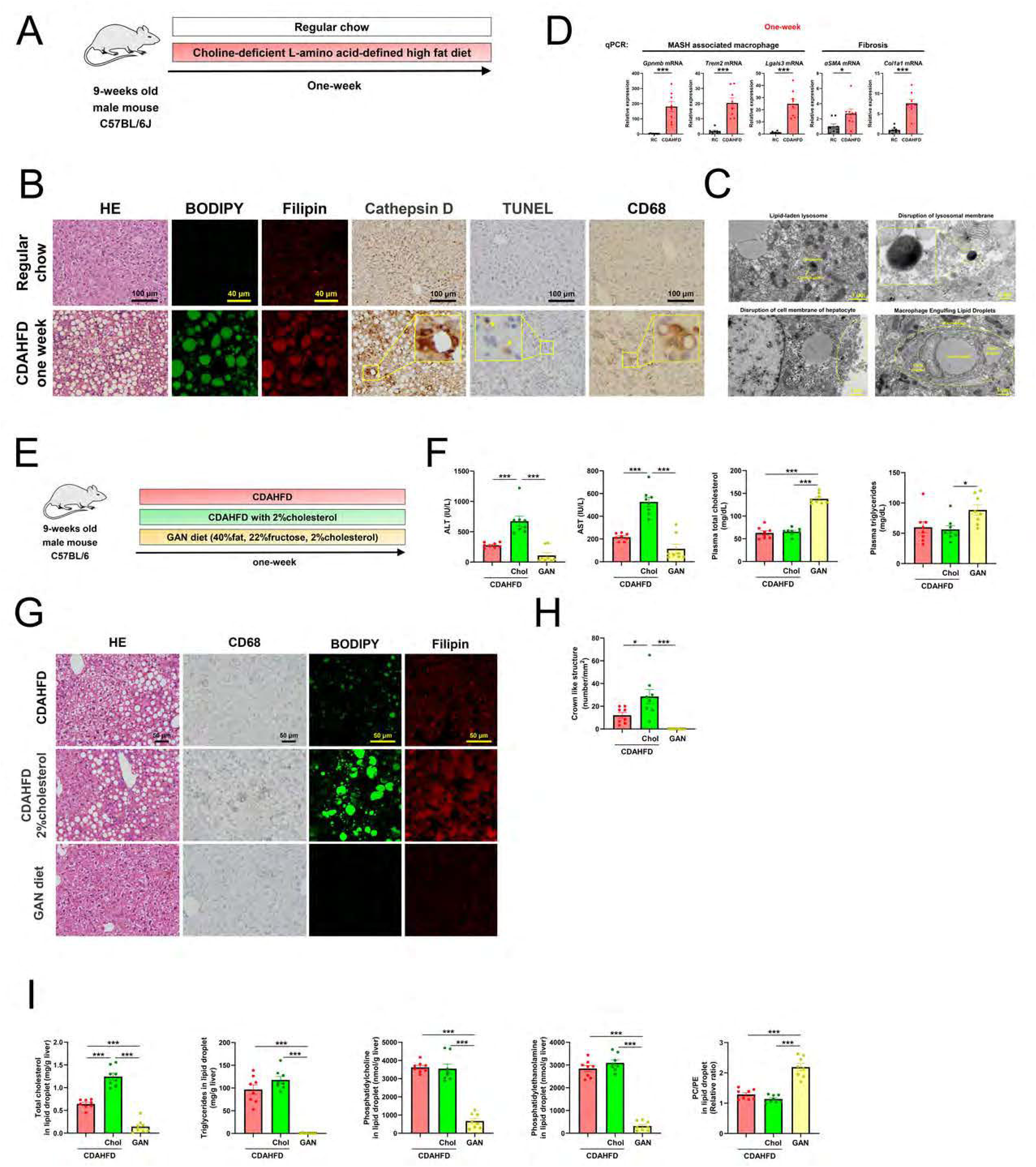
Choline-deficient L-amino acid-defined high-fat diet induces steatosis via enhanced hepatic uptake of chylomicron remnants. (A) Study design. C57BL/6J mice were divided into regular chow (RC)-fed mice or choline-deficient, L-amino acid-defined, high-fat diet (CDAHFD)-fed mice for one week. (B) Liver sections were stained with hematoxylin, eosin (HE), BODIPY, Filipin, Cathepsin D, Terminal deoxynucleotidyl transferase dUTP nick-end labeling (TUNEL) and CD68. CDAHFD induced macrovesicular steatosis. BODIPY and Filipin staining demonstrated that free cholesterol was included in liver lipid droplets. Leakage of cathepsin D into cytoplasm indicated disruption of the lysosomal membrane. TUNEL staining revealed dead hepatocytes. CD68 staining revealed crown-like structures. (C) Representative transmission electron microscopy images of one week CDAHFD-fed mice’s liver. Transmission electron microscopy demonstrated lipid-laden lysosome, disruption of lysosomal membrane, disruption of cell membrane of hepatocyte and macrophage engulfing lipid droplets. (D) Quantitative real-time polymerase chain reaction (RT-qPCR) analysis of liver tissues. CDAHFD one week feeding increased mRNA expression of metabolic dysfunction-associated steatohepatitis (MASH) associated macrophage markers (Gpnmb, Trem2, and Lgals3), and fibrosis markers (αSMA, and Col1a1). (E) Study design. C57BL/6J mice were divided into a choline-deficient, L-amino acid-defined, high-fat diet (CDAHFD)-fed mice, CDAHFD with 2% cholesterol-fed mice, Gubra-Amylin NASH (GAN) diet-fed mice. (F) Plasma ALT, AST, total cholesterol and triglycerides levels. CDAHFD with 2% cholesterol-fed mice showed increased ALT and AST levels compared to CDAHFD-fed mice and GAN-diet fed mice. GAN diet-fed mice showed increased plasma total cholesterol and triglyceride levels compared to CDAHFD with 2% cholesterol-fed mice. (G) Liver sections were stained with hematoxylin, eosin (HE), CD68, BODIPY, and Filipin. CDAHFD and CDAHFD with 2% cholesterol feeding for one week induced steatosis. GAN diet feeding for one week did not cause apparent steatosis. (H) The number of crown-like structures in the liver increased in CDAHFD with 2% cholesterol-fed mice compared to CDAHFD-fed mice and GAN diet-fed mice. (I) The metabolites levels in liver lipid droplet. Total cholesterol and triglycerides increased in CDAHFD-fed mice and CDAHFD with 2% cholesterol-fed mice compared to GAN diet-fed mice. In addition, total cholesterol in CDAHFD with 2% cholesterol-fed mice was higher than that of CDAHFD-fed mice. The level of phosphatidylcholine and phosphatidylethanolamine increased in CDAHFD-fed mice and CDAHFD with 2% cholesterol-fed mice compared to GAN diet-fed mice. The ratio of phosphatidylcholine (PC) to phosphatidylethanolamine (PE) decreased in CDAHFD-fed mice and CDAHFD with 2% cholesterol-fed mice compared to GAN diet-fed mice. Data are presented as mean ± SEM. Groups were compared by one-way ANOVA followed by Tukey’s multiple comparisons test.

CDAHFD induced macrovesicular hepatic steatosis (Figure 2B). BODIPY and Filipin staining demonstrated that free cholesterol was included in liver lipid droplets. Leakage of cathepsin D into cytoplasm indicated disruption of the lysosomal membrane. Lysosomal rupture with cathepsin release causes various types of cell death (15) and consistent with these findings TUNEL staining revealed apoptotic hepatocytes. CD68 staining revealed crown-like structures. Transmission electron microscopy demonstrated lipid-laden lysosome, disruption of lysosomal membrane, disruption of cell membrane of hepatocyte and macrophage engulfing lipid droplets (Figure 2C).

We next measured mRNA expression of genes associated with MASH in this one-week CDAHFD model. Hepatic macrophages are thought to play key roles in the pathogenesis of MASH. Resident Kupffer cells are replaced by distinct subsets of bone marrow derived macrophages (16). Previously, experiments using single-cell RNA sequencing identified MASH-associated macrophages that were associated with progression to fibrosis (17). Additional studies suggested that *Trem2^High^-Gpnmb^High^* macrophages express *Lgals3* (*18, 19*), a cytokine that promotes hepatic stellate cells activation (20). We observed that one week feeding of the CDAHFD diet induced expression of *Gpnmb*, *Trem2* and *Lgals3*. These changes were associated with increased mRNA expression of fibrosis markers (*αSMA*, and *Col1a1*) (Figure 2D). Furthermore, the profile of mRNA changes that we observed in this one-week study was similar to that obtained from the livers of 12-week CDAHFD-fed mice [Gene Expression Omnibus data set (GSE120977)] (Figure S2B). Thus, the early changes in gene expression after 1 week appear to be sustained with chronic CDAHFD feeding.

### Cholesterol supplementation induces cholesterol in liver lipid droplets, especially under choline deficiency

To test our hypothesis that hepatic cholesterol accumulation in lipid droplets promotes MASH in additional mouse models of MASH, C57BL/6J mice were divided into CDAHFD-fed mice, CDAHFD with 2% cholesterol-fed mice (CDAHFD+Chol), Gubra-Amylin (GAN) diet-fed mice (Figure 2E). The GAN diet includes 40% kcal fat (palm oil, soybean oil), 22% wt fructose, sufficient choline and 2% cholesterol. We selected a CDAHFD containing 2% cholesterol to compare with the GAN diet, which also includes 2% cholesterol. CDAHFD+Chol fed mice had an exacerbation of the MASH phenotype as reflected by higher plasma ALT and AST levels compared to CDAHFD-fed mice and GAN-diet-fed mice (Figure 2F). GAN diet-fed mice showed increased plasma total cholesterol and triglyceride levels compared to CDAHFD with 2% cholesterol-fed mice (Figure 2F). Interestingly while CDAHFD (with or without Chol) feeding for one week resulted in hepatic steatosis, GAN diet feeding for one week did not cause hepatic steatosis (Figure 2G). Consistent with the increased hepatic inflammation, CDAHFD+Chol feeding promoted further increases in cholesterol content in liver lipid droplets. The number of crown-like structures in the liver was highest in the CDAHFD + Chol-fed mice compared to CDAHFD-fed mice and GAN diet-fed mice (Figure 2H). Total cholesterol in liver lipid droplets was highest in the CDAHFD+Chol mice compared to both CDAHFD alone and GAN diet (Figure 2I).

In contrast, liver lipid droplet triglyceride increased to a similar extent in both CDAHFD-fed mice and CDAHFD + Chol-fed mice compared to GAN diet-fed mice (Figure 2I). Similarly, the concentration of PC and PE in liver lipid droplets increased in both CDAHFD and CDAHFD+Chol-fed mice compared to GAN diet-fed mice. The ratio of PC to PE decreased in both CDAHFD and CDAHFD+Chol-fed mice compared to GAN diet-fed mice. Taken together these data demonstrate that, dietary cholesterol supplementation in addition to a choline-deficient diet profoundly increased cholesterol in liver lipid droplets compared to cholesterol supplementation on choline sufficient diet conditions. This is consistent with the fact that the GAN diet induces MASH more slowly compared to CDAHFD (8). The close association between liver lipid droplet cholesterol and worsening histological changes further points to the accumulation of lipid droplet cholesterol content as being more closely tied to the development of MASH than other putative factors, such as dietary cholesterol content, plasma cholesterol concentration or intrahepatic PC to PE ratio.

### Bempedoic acid alleviates liver inflammation and fibrosis due to choline-deficient L-amino acid-defined high-fat diet via decreasing cholesterol content in liver lipid droplets

To test our hypothesis that CDAHFD leads to MASH by increasing cholesterol content in liver lipid droplets, we treated CDAHFD-fed mice with bempedoic acid (Figure 3A). Bempedoic acid works as an ATP-citrate lyase inhibitor, which is clinically used as a plasma cholesterol-lowering drug (21). While bempedoic acid has been shown to be effective in reducing hepatic steatosis in previous studies (22–24), its effect on cholesterol content in liver lipid droplets has not yet been studied. Bempedoic acid treatment reduced plasma ALT and AST (Figure 3B), hepatic steatosis (Figure 3C) and the number of crown-like structures compared to the control mice (Figure 3C and 3D). BODIPY and Filipin staining demonstrated decreased free cholesterol in liver lipid droplets of bempedoic acid-treated mice (Figure 3C). Bempedoic acid treatment also reduced triglycerides and total cholesterol content in liver lipid droplets (Figure 3E). The level of PC in liver lipid droplets decreased in bempedoic acid-treated mice whereas PE and the ratio of PC to PE remained unchanged. Consistent with pathological findings (Figure 3C), RT-qPCR demonstrated a decreased mRNA expression of MASH-associated macrophage markers (*Gpnmb, Trem2, and Lgals3*), and fibrosis marker (*Col1a1*) in bempedoic acid-treated mice liver (Figure 3F). Taken together, these data demonstrate that decreasing hepatic cholesterol synthesis by inhibiting ATP-citrate lyase reduces hepatic lipid droplet cholesterol (and triglyceride) content and attenuates the MASH phenotype in CDAHFD-fed mice.

**Figure 3.**
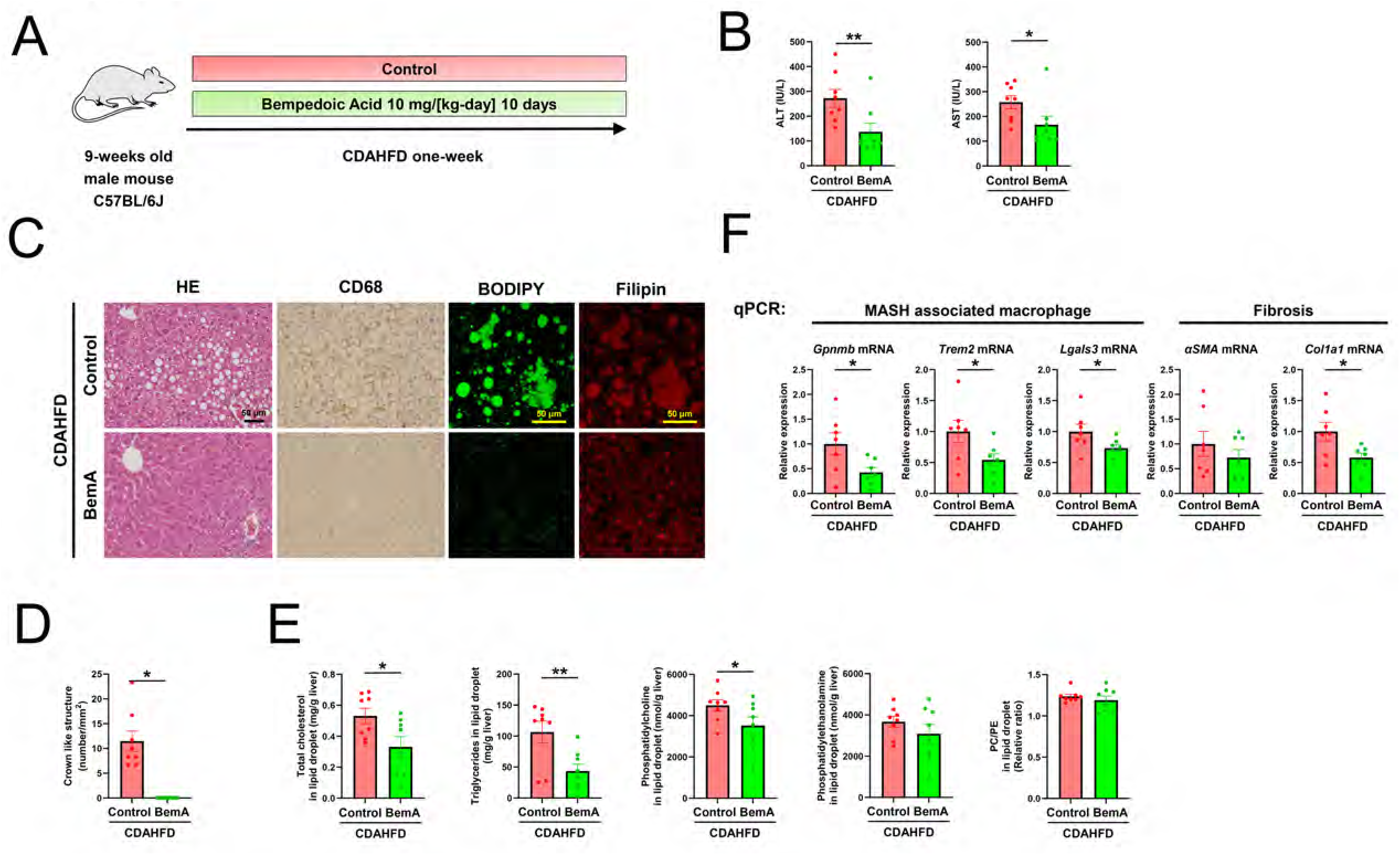
Bempedoic acid alleviates liver inflammation and fibrosis due to choline-deficient L-amino acid-defined high-fat diet via decreasing cholesterol content in liver lipid droplets. (A) Study design. C57BL/6J mice were divided into a control group and bempedoic acid-treated group. Both groups were fed a CDAHFD for one week. (B) ALT and AST in bempedoic acid (BemA) treated mice decreased compared to control mice. (C) Liver sections were stained with hematoxylin, eosin (HE), CD68, BODIPY, and Filipin. BemA treatment prevented steatosis. CD68 staining revealed decreased crown-like structures. BODIPY and Filipin staining demonstrated decreased free cholesterol in liver lipid droplet. (D) The number of crown-like structures decreased in BemA-treated mice liver. (E) The metabolites levels in liver lipid droplet. Total cholesterol and triglycerides decreased in BemA-treated mice. The level of phosphatidylcholine (PC) decreased in BemA-treated mice. Phosphatidylethanolamine (PE) did not show a difference. The ratio of PC to PE did not show a difference. (F) Quantitative real-time polymerase chain reaction (RT-qPCR) analysis of liver tissues demonstrated a decreased mRNA expression of MASH-associated macrophage markers (*Gpnmb*, *Trem2*, and *Lgals3*), and fibrosis marker (*Col1a1*) in BemA-treated mice. Data are presented as mean ± SEM. Groups were compared by Unpaired one-sided Student’s t-test.

### Atorvastatin alleviates liver inflammation and fibrosis due to choline-deficient L-amino acid-defined high-fat diet via decreasing cholesterol content in liver lipid droplets

To further examine the role of liver lipid droplet cholesterol on MASH pheonotype in CDAHFD-fed mice, we evaluated whether short-term atorvastatin (statin) treatment would decrease cholesterol content in liver lipid droplets and alleviate MASH in the CDAHFD one-week model (Figure S3A). Statin treatment resulted in reductions of plasma ALT, AST (Figure S3B) and reduced hepatic steatosis (Figure S3C). There was a reduction in the number of crown-like structures compared to the control mice (Figure S3C and S3D). BODIPY and Filipin staining demonstrated decreased free cholesterol in liver lipid droplets of statin-treated mice (Figure S3C). Consistent with BODIPY and Filipin staining, statin treatment also reduced triglycerides and total cholesterol content in liver lipid droplets (Figure S3E). The level of PC and PE in liver lipid droplets did not differ between control and statin-treated mice (Figure S3E). As a result, the ratio of PC to PE in liver lipid droplets did not show a difference. Consistent with pathological findings (Figure S3C), RT-qPCR demonstrated a decreased mRNA expression of MASH-associated macrophage markers (*Gpnmb, Trem2, and Lgals3*), and fibrosis markers (*αSMA* and *Col1a1*) in statin-treated mice liver (Figure S3F).

### Coenzyme A synthase knockdown alleviates liver inflammation and fibrosis due to Choline-deficient L-amino acid-defined high-fat diet via decreasing cholesterol in liver lipid droplets

Coenzyme A synthase (COASY) plays a crucial role in coenzyme A (CoA) production from 4’-phosphopantetheine. COASY generates CoA which is required for the synthesis of acetyl CoA, a key substrate for the cholesterol synthetic pathway (Figure 4A). To evaluate whether Coasy knockdown decreases cholesterol in liver lipid droplets and alleviates MASH in the CDAHFD one-week model, we divided C57BL6 mice into a GalNAc control ASO-treated mouse group and a GalNAc *Coasy* ASO-treated mouse group. Both groups were subjected to a CDAHFD feeding for one week (Figure 4B).

**Figure 4.**
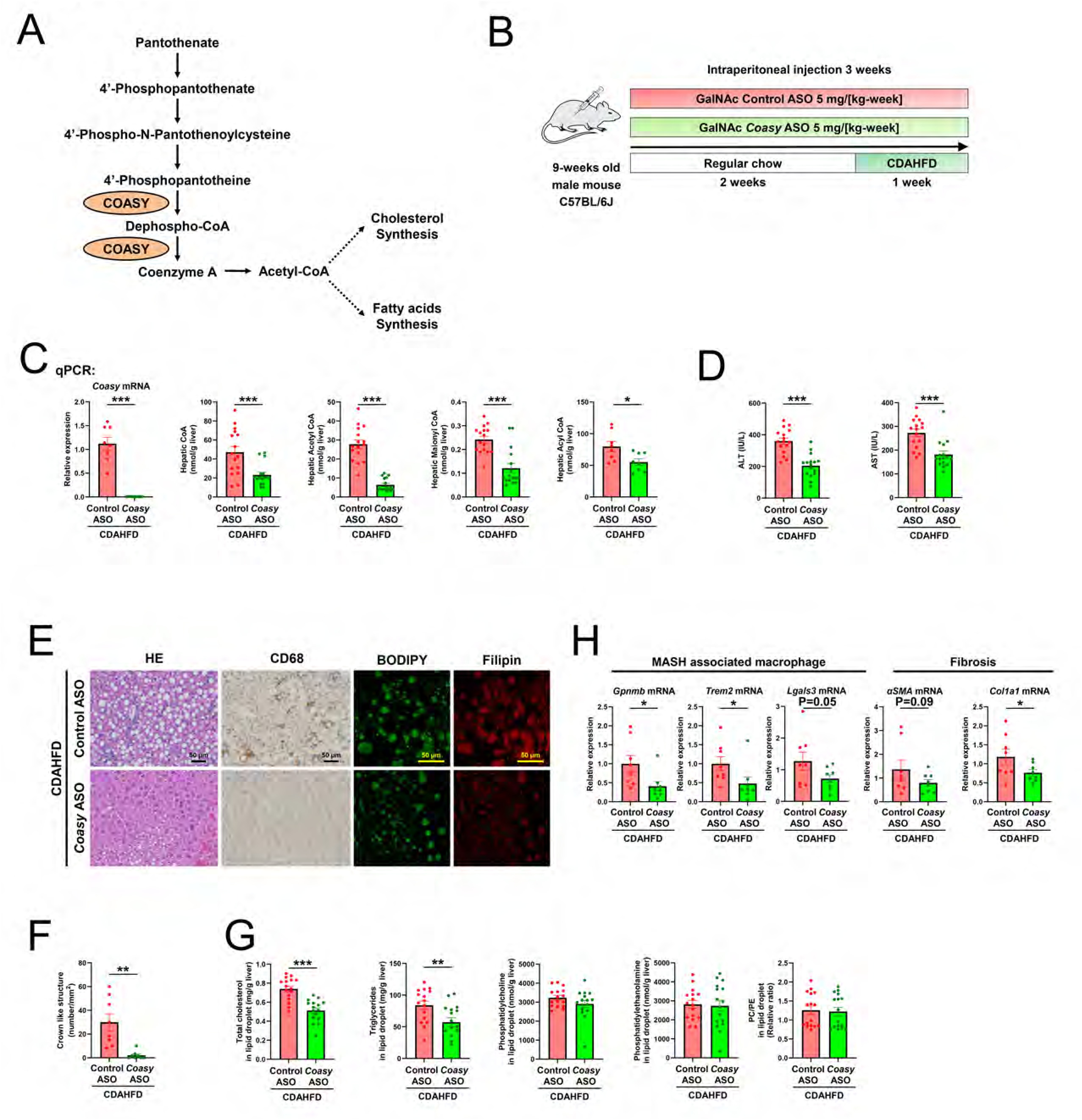
Coenzyme A synthase knockdown alleviates liver inflammation and fibrosis due to Choline-deficient L-amino acid-defined high-fat diet via decreasing cholesterol in liver lipid droplets. (A) Coenzyme A biosynthesis pathway. Coenzyme A synthase (COASY) converts 4’-Phosphopantetheine into Coenzyme A (CoA) by adenylation and phosphorylation. CoA subsequently catalyzes the synthesis of acetyl CoA. Acetyl CoA functions as a substrate throughout the cholesterol synthesis or fatty acids synthesis pathway. (B) Study design. C57BL/6J mice were divided into GalNAc control ASO-treated mice and GalNAc Coasy ASO-treated mice. Both groups were fed a choline-deficient, L-amino acid-defined, high-fat diet (CDAHFD) for one week. (C) Quantitative real-time polymerase chain reaction (RT-qPCR) analysis of liver tissues confirmed the knockdown effect of Coasy ASO. Consistently, Coasy ASO treatment decreased hepatic Coenzyme A species, including CoA, Acetyl CoA, Malonyl CoA, and Acyl CoA. (D) ALT and AST in Coasy ASO-treated mice decreased compared to control ASO-treated mice. (E) Liver sections were stained with hematoxylin, eosin (HE), CD68, BODIPY, and Filipin. Coasy ASO treatment alleviated steatosis. Accordingly, CD68 staining revealed decreased crown-like structures. BODIPY and Filipin staining demonstrated decreased free cholesterol in liver lipid droplets. (F) The number of crown-like structures decreased in Coasy ASO-treated mice liver. (G) The metabolites levels in liver lipid droplet. Total cholesterol and triglycerides decreased in Coasy ASO-treated mice. The level of phosphatidylcholine (PC) and phosphatidylethanolamine (PE) decreased in Coasy ASO-treated mice. The ratio of PC to PE remained unchanged. (H) Quantitative real-time polymerase chain reaction (RT-qPCR) analysis of liver tissues demonstrated a decreased mRNA expression of MASH-associated macrophage markers (Gpnmb,Trem2 and Lgals3) and fibrosis markers (αSMA, and Col1a1) in Coasy ASO-treated mice. Data are presented as mean ± SEM. Groups were compared by Unpaired one-sided Student’s t-test.

*Coasy* mRNA expression was decreased by >95% with *Coasy* ASO, confirming the efficacy of *Coasy* ASO (Figure 4C). *Coasy* ASO treatment decreased all hepatic Coenzyme A species, including CoA, acetyl CoA, malonyl CoA, and acyl CoA (Figure 4C). Plasma ALT and AST concentrations decreased in *Coasy* ASO-treated mice compared to control ASO-treated mice (Figure 4D). HE and BODIPY staining demonstrated that Coasy knockdown decreased liver lipid droplets (Figure 4E). The number of crown-like structures decreased in *Cosasy* ASO-treated mouse livers (Figures 4E and 4F). BODIPY and Filipin staining demonstrated decreased free cholesterol in liver lipid droplets of *Coasy* ASO-treated mice (Figure 4E). Consistent with BODIPY and Filipin staining, triglycerides and total cholesterol in liver lipid droplets decreased in *Coasy* ASO-treated mice (Figure 4G). However, neither the concentration of PC and PE in liver lipid droplets nor the PC: PE ratio changed with *Coasy* ASO treatment (Figure 4G). Consistent with pathological findings, RT-qPCR demonstrated decreased mRNA expression of MASH-associated macrophage markers (*Gpnmb*, *Trem2, and Lgals3*), and fibrosis markers (*αSMA*, and *Col1a1*) in *Coasy* ASO treated mice liver (Figure 4H).

It is generally accepted that there is no mouse model of MASH that manifests all of the characteristics of human MASH. To address this concern we also examined the effect of Coasy knockdown on MASH pathogenesis in a prolonged (40 weeks) GAN-diet fed mouse model of MASH (Figure S4A). Plasma ALT and AST concentrations decreased in Coasy ASO-treated mice compared to control ASO-treated mice under GAN-diet feeding (Figure S4B). The decrease in liver fibrosis in Coasy ASO-treated mice was confirmed by liver hydroxyproline assay (Figure S4B). Total cholesterol, triglycerides, PC, and PE in liver lipid droplets decreased in Coasy ASO-treated mice (Figure S4C). The PC: PE ratio increased with Coasy ASO treatment. Taken together with our prior results demonstrating a protective effect of Coasy ASO treatment in the development of MASH in a CDAHFD mouse model of MASH these results confirm the protective effects of Coasy knockdown in liver on the progression of MASH in a well-established prolonged GAN diet fed mouse model of MASH.

### Cholesterol supplementation negates the protective effect of Coasy ASO on a choline-deficient L-amino acid-defined high-fat diet one-week mice model

To test whether increases in cholesterol per se could negate the protective effect of *Coasy* knockdown, *Coasy* ASO-treated C57BL/6J mice were divided into mice fed either CDAHFD alone or CDAHFD with 2% cholesterol (Figure 5A). As before, CDAHFD+Chol exacerbated the MASH phenotype with higher transaminase concentrations (Figure 5B), histological evidence of increased cholesterol in liver lipid droplets (Figure 5C) and increased number of crown like structures (Figures 5C and 5D). Consistent with these findings, triglycerides and total cholesterol in liver lipid droplets increased in cholesterol-supplemented mice (Figure 5E). In addition, there was a modest increase in lipid droplets PC and PE in CDAHFD+Chol-fed mice (Figure 5E). The ratio of PC to PE in liver lipid droplets was not different between the two diet groups. RT-qPCR demonstrated an increased mRNA expression of MASH-associated macrophage markers (*Gpnmb*, *Trem2, and Lgals3) and fibrosis markers (αSMA* and *Col1a1*) in cholesterol-supplemented mice liver (Figure 5F). Thus cholesterol supplementation abrogated the protective effects of *Coasy* ASO in CDAHFD-fed mice.

**Figure 5.**
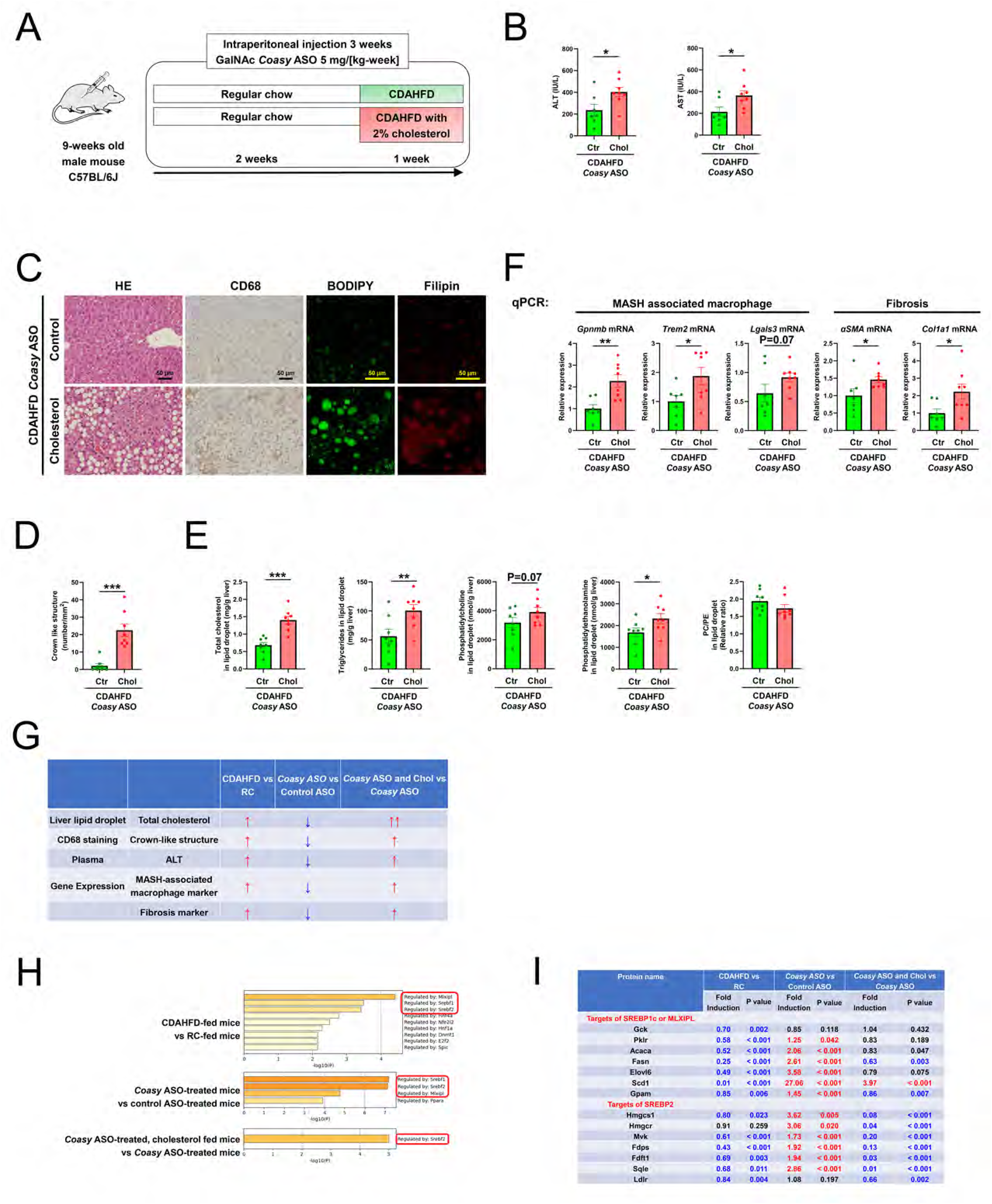
Cholesterol supplementation negates the protective effect of Coasy ASO on a Choline-deficient L-amino acid-defined high-fat diet one-week mice model. (A) Study design. C57BL/6J mice were divided into choline-deficient, L-amino acid-defined, high-fat diet (CDAHFD) treated mice and CDAHFD with 2 % cholesterol treated mice. Both groups were injected with GalNAc *Coasy* ASO. (B) Cholesterol (Chol) dietary supplementation increased plasma ALT and AST levels compared to control (Ctr) mice. (C) Liver sections were stained with hematoxylin, eosin (HE), CD68, BODIPY, and Filipin. Dietary cholesterol supplementation induced steatosis. CD68 staining revealed increased crown-like structures. BODIPY and Filipin staining demonstrated increased free cholesterol in liver lipid droplet. (D) The number of crown-like structures increased in dietary cholesterol-supplemented mice liver. (E) The metabolites levels in liver lipid droplet. Total cholesterol and triglycerides increased in cholesterol-overload mice. The level of phosphatidylcholine (PC) and phosphatidylethanolamine increased (PE) in cholesterol-supplemented mice. The ratio of PC to PE remained unchanged. (F) Quantitative real-time polymerase chain reaction (RT-qPCR) analysis of liver tissues showed an elevated mRNA expression of MASH-associated macrophage markers (*Gpnmb, Trem2 and Lgals3*) as well as fibrosis markers (*αSMA, and Col1a1*) in cholesterol-overload mice. Data are presented as mean ± SEM. Groups were compared by Unpaired one-sided Student’s t-test. (G) Summary of four studies of one-week CDAHFD studies. (H) Enrichment analysis of comprehensive proteome analysis. Three comparisons were made: (1) CDAHFD-fed mice (n=5) vs. Regular chow (RC)-fed mice (n=5); (2) *Coasy* ASO-CDAHFD-treated mice (n=5) vs. Control ASO-CDAHFD-treated mice (n=5); and (3) *Coasy* ASO-CDAHFD with 2% cholesterol-treated mice (n=5) vs. *Coasy* ASO-CDAHFD-treated mice (n=5). (I) Hepatic protein expression of targets of SREBP1c, MLXIPL, and SREBP2

### Comprehensive proteome analysis revealed that choline-deficient L-amino acid-defined high-fat diet compensatory suppress targets of SREBP1c and SREBP2, and Coasy knockdown reverses this alteration

Together, these data support the hypothesis that cholesterol accumulation in liver lipid droplets plays a critical role in the rapid development of MASH in CDAHFD-fed mice (Figure 5G). We next performed a comprehensive proteome analysis and then subjected it to enrichment analysis to identify potential pathways that may be altered in each experimental group compared to the control group. We found that Mlxipl, Srebf1, and Srebf2 pathways were significantly altered in CDAHFD-fed mice vs. RC-fed mice and *Coasy* ASO-treated mice vs. control ASO-treated mice (Figure 5H). The Srebf2 pathway was the most enriched in *Coasy* ASO-treated cholesterol-fed mice vs. *Coasy* ASO-treated mice. Hepatic protein expression of targets of MLXIPL, SREBP1c and SREBP2 were suppressed in CDAHFD-fed mice compared to RC-fed mice and activated in *Coasy* ASO-treated mice compared to control ASO-treated mice (Figure 5I). The expression of SREBP2 targets was increased with *Coasy* ASO treatment but suppressed in *Coasy* ASO and cholesterol-fed mice.

Treatments that decreased intracellular cholesterol increased the expression of SREBP2 targets which likely represents a compensatory response to cholesterol depletion in *Coasy* ASO-treated mice. Interestingly, though *Coasy* ASO also increased a number of SREBP1c targets, perhaps to compensate for decreased fatty acid synthesis, cholesterol supplementation appeared to prevent upregulation of SREBP1c targets, possibly representing a degree of crosstalk between cholesterol and fatty acid biosynthetic pathways.

### Hydroxysteroid 17-dehydrogenase 13 knockdown prevents liver inflammation and fibrosis due to choline-deficient L-amino acid-defined high-fat diet via decreasing cholesterol in liver lipid droplets

Recently, GWAS identified 17 loci associated with MASLD (25), including several genetically validated potential targets for treating MASH (26, 27). Patatin-like phospholipase domain-containing protein 3 (PNPLA3) I148M increases the risk (28). The hydroxysteroid 17-beta dehydrogenase 13 (HSD17B13) splice site variant (rs72613567) decreases the risk of MASH (29). Among variants associated with risk of MASLD/MASH, the effect size of PNPLA3 and HSD17B13 variants are larger than other variants (27). In addtion, the HSD17B13 splice site variant (rs72613567) is found in 18% of the global population (30, 31). While HSD17B13 is a lipid droplet surface protein, it has not been shown to directly modulate the cholesterol content of lipid droplets. The loss of function variant, rs72613567:TA, in the HSD17B13 gene confers protection against both alcoholic and metabolic liver diseases(29). To evaluate whether Hsd17b13 knockdown decreases cholesterol in liver lipid droplets and alleviates MASH in the CDAHFD one-week model, we divided C57BL/6J mice into GalNAc control ASO treated mice, and GalNAc *Hsd17b13* ASO-treated mice under CDAHFD one week (Figure 6A). RT-qPCR analysis of liver tissues confirmed the knockdown effect of *Hsd17b13* ASO (Figure 6B). *Hsd17b13* ASO decreased plasma AST and ALTcompared to the control ASO (Figure 6C). HE staining demonstrated that Hsd17b13 knockdown did not significantly affect steatosis (Figure 6D).

**Figure 6.**
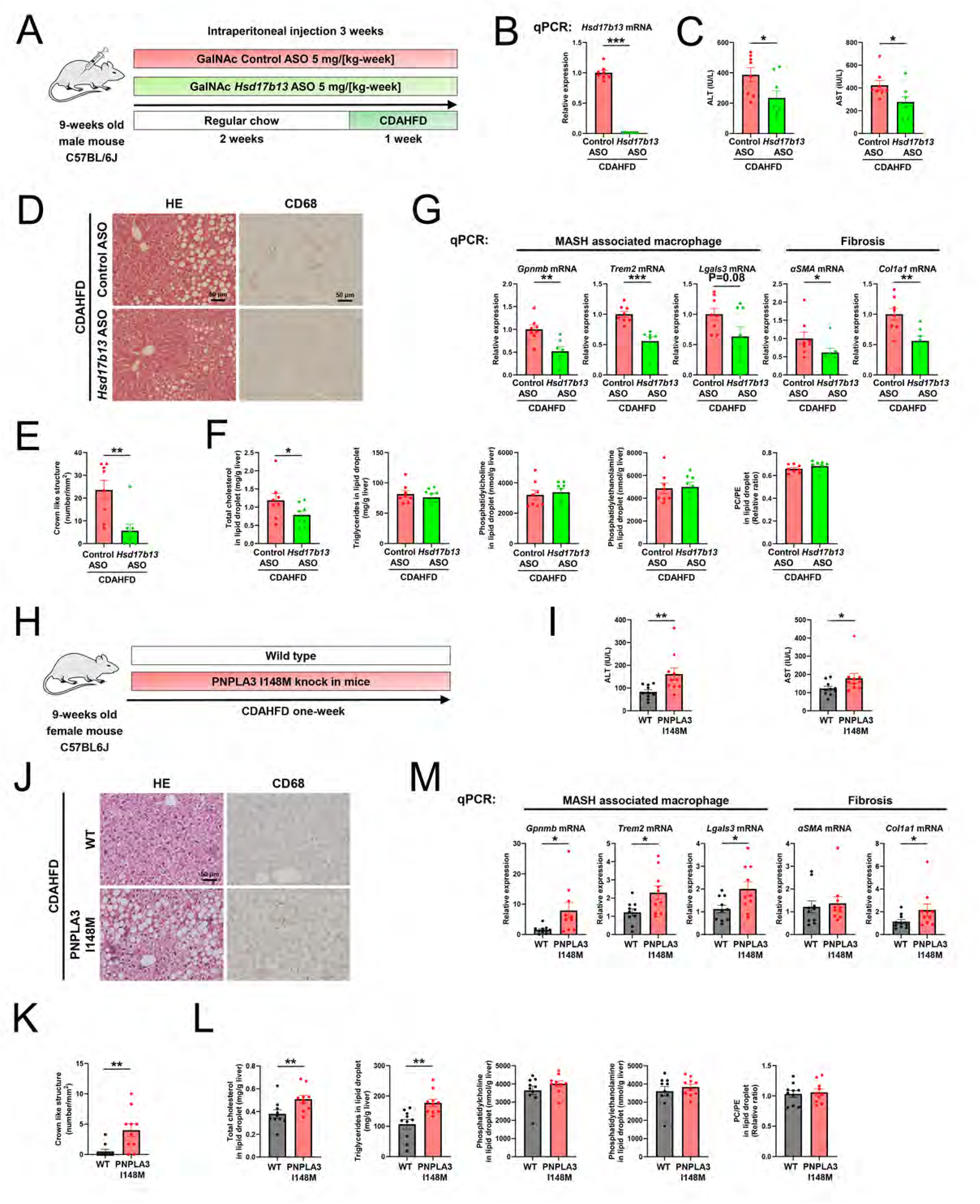
Hydroxysteroid 17-beta dehydrogenase 13 knockdown alleviates MASH due to CDAHFD via decreasing cholesterol in liver lipid droplets and Patatin-like phospholipase domain-containing protein 3 I148M knock-in aggravates MASH due to CDAHFD via increasing in liver lipid droplets. (A) Study design. C57BL/6J mice were divided into GalNAc control ASO-treated mice and GalNAc Hydroxysteroid 17-beta dehydrogenase 13 (Hsd17b13) ASO-treated mice. Both groups were fed a choline-deficient, L-amino acid-defined, high-fat diet (CDAHFD) for one week. (B) Quantitative real-time polymerase chain reaction (RT-qPCR) analysis of liver tissues confirmed the knockdown effect of Hsd17b13 ASO. (C) Plasma ALT and AST levels in Hsd17b13 ASO-treated mice decreased compared to control mice. (D) Liver sections were stained with hematoxylin, eosin (HE) and CD68. Hsd17b13 ASO did not significantly affect steatosis. CD68 staining revealed decreased crown-like structures. (E) The number of crown-like structures decreased in Hsd17b13 ASO-treated mice liver. (F) The metabolites levels in liver lipid droplet. Total cholesterol decreased in Hsd17b13 ASO-treated mice. The level of triglycerides, phosphatidylcholine (PC) and phosphatidylethanolamine (PE) did not change. The ratio of PC to PE did not change. (G) Quantitative real-time polymerase chain reaction (RT-qPCR) analysis of liver tissues demonstrated a decreased mRNA expression of MASH-associated macrophage markers (Gpnmb, Trem2 and Lgals3) and fibrosis markers (αSMA and Col1a1) in Hsd17b13 ASO-treated mice. (H) Study design. Patatin-like phospholipase domain-containing protein 3 (Pnpla3) I148M knock-in mice and wild-type mice were fed with a choline-deficient, L-amino acid-defined, high-fat diet (CDAHFD) for one week. (I) ALT and AST in Pnpla3 I148M knock-in mice increased compared to wild-type mice. (J) Liver sections were stained with hematoxylin, eosin (HE), and CD68. Pnpla3 I148M showed macrovesicular steatosis compared to wild-type mice. CD68 staining revealed increased crown-like structures. (K) The number of crown-like structures increased in Pnpla3 I148M knock-in mice liver. (L) The metabolite levels in the liver lipid droplet. Total cholesterol and triglycerides increased in Pnpla3 I148M knock-in mice. The level of phosphatidylcholine (PC) and phosphatidylethanolamine (PE) did not change. The ratio of PC to PE did not change. (M) Quantitative real-time polymerase chain reaction (RT-qPCR) analysis of liver tissues demonstrated an increased mRNA expression of MASH-associated macrophage markers (Gpnmb, Trem2, and Lgals3), and fibrosis markers (αSMA, and Col1a1) in Pnpla3 I148M knock-in mice. Data are presented as mean ± SEM. Groups were compared by Unpaired one-sided Student’s t-test.

The number of crown-like structures decreased in *Hsd17b13* ASO-treated mice’s livers (Figures 6D and 6E). Total cholesterol in liver lipid droplets decreased in *Hsd17b13* ASO-treated mice, while the level of triglycerides, PC and PE in liver lipid droplets did not differ between control and *Hsd17b13* ASO-treated mice (Figure 6F). As a result, the ratio of PC to PE in liver lipid droplets did not show a difference. Consistent with pathological findings, RT-qPCR demonstrated a reduced mRNA expression of MASH-associated macrophage markers (*Gpnmb, Trem2, and Lgals3*) and fibrosis markers (*αSMA* and *Col1a1*) in *Hsd17b13* ASO-treated mice (Figure 6G).

### Cholesterol supplementation negates the protective effect of Hsd17b13 ASO on a choline-deficient L-amino acid-defined high-fat diet one-week mice model

To test whether increases in cholesterol per se could negate the protective effect of *Hsd17b13* knockdown, *Hsd* ASO-treated C57BL/6J mice were divided into mice fed either CDAHFD alone or CDAHFD with 2% cholesterol (Figure S5A). As before, CDAHFD+Chol exacerbated the MASH phenotype with higher transaminase concentrations (Figure S5B), and increased number of crown like structures (Figures S5C and S5D). Consistent with these findings, triglycerides and total cholesterol in liver lipid droplets increased in cholesterol-supplemented mice (Figure S5E). There was no change in lipid droplets PC and PE in CDAHFD+Chol-fed mice (Figure S5E). The ratio of PC to PE in liver lipid droplets was not different between the two diet groups. RT-qPCR demonstrated an increased *Gpnmb*, *Trem2 and Col1a1* mRNA expression in cholesterol-supplemented mice liver (Figure S5F). Thus cholesterol supplementation abrogated the protective effects of *Hsd17b13* ASO in CDAHFD-fed mice.

### Patatin-like phospholipase domain-containing protein 3 I148M knock-in mice increase cholesterol in liver lipid droplets

Next we investigated the role of PNPLA3 I148M in the development of MASH. One possible mechanism is that PNPLA3 148M competes with ATGL for the interaction with CGI-58, reducing intrahepatic lipolysis (32). To evaluate whether PNPLA3 I148M increases cholesterol content in liver lipid droplets, we studied PNPLA3 I148M knock-in mice and wild-type mice under CDAHFD one-week treatment (Figure 6H). Plasma ALT and AST in PNPLA3 I148M mice were higher than in wild-type mice (Figure 6I). The number of crown-like structures increased in the liver of PNPLA3 I148M mice (Figure 6J and 6K). Total cholesterol and triglycerides in liver lipid droplets increased in PNPLA3 I148M mice (Figure 6L). The level of PC and PE in liver lipid droplets did not differ between wild-type and PNPLA3 I148M mice (Figure 6L). As a result, the ratio of PC to PE in liver lipid droplets did not differ. Consistent with pathological findings, RT-qPCR demonstrated an increased mRNA expression of MASH-associated macrophage markers (*Gpnmb, Trem2,* and *Lgals3*), and fibrosis marker (*Col1a1*) in PNPLA3 I148M mice (Figure 6M).

We next evaluated whether bempedoic acid treatment would decrease cholesterol in liver lipid droplets and alleviate MASH in PNPLA3 I148M knock-in mice fed with CDAHFD (Figure S6A). Bempedoic acid treatment reduced plasma ALT and AST (Figure S6B), hepatic steatosis (Figure S6C), and the number of crown-like structures compared to the control mice (Figure S6C and S6D). Bempedoic acid treatment also reduced triglycerides and total cholesterol content in liver lipid droplets (Figure S6E). PC and PE in liver lipid droplets did not show a difference. The ratio of PC to PE did not show a difference. RT-qPCR demonstrated a decreased mRNA expression of MASH-associated macrophage markers (Gpnmb, Trem2, and Lgals3), and fibrosis markers (αSMA and Col1a1) in bempedoic acid-treated mice liver (Figure S6F). Taken together, these data demonstrate that decreasing hepatic cholesterol synthesis by bempedoic acid reduces hepatic lipid droplet cholesterol (and triglyceride) content and attenuates the MASH phenotype in PNPLA3 I148M knock-in mice fed with CDAHFD.

### Lipid droplet cholesterol is a key factor in human MASLD progression associated with variants of PNPLA3 and HSD17B13

To further validate the key role of liver lipid droplet cholesterol in human MASH, we evaluated metabolites in lipid droplets of human liver samples. Human liver samples were divided into five groups (no MASL, MASL without fibrosis, MASH without fibrosis, MASH and F>0, MASL and F>0 but no MASH) (Table S1). Total cholesterol and triglycerides in liver lipid droplets were higher in MASH and F>0, compared to other groups (Figure 7A). Next, we evaluated these samples from the based on the relatively common PNPLA3 I148M variant and the HSD17B13 splice variant (rs72613567), which have been shown to be aggravating or protecting variants for the development of MASH, respectively (Table S2). We compared the PNPLA3 I148M variant group without HSD17B13 splice site variants as the high genetic risk group to the HSD17B13 splice site variant group without PNPLA3 I148M variants as the low genetic risk group (Figure 7B). Total cholesterol, triglycerides PC and PE in liver lipid droplets were higher in PNPLA3 I148M variants without HSD17B13 splice site variants than in HSD17B13 splice site variants without PNPLA3 I148M variants. The ratio of PC to PE in liver lipid droplets did not differ.

**Figure 7.**
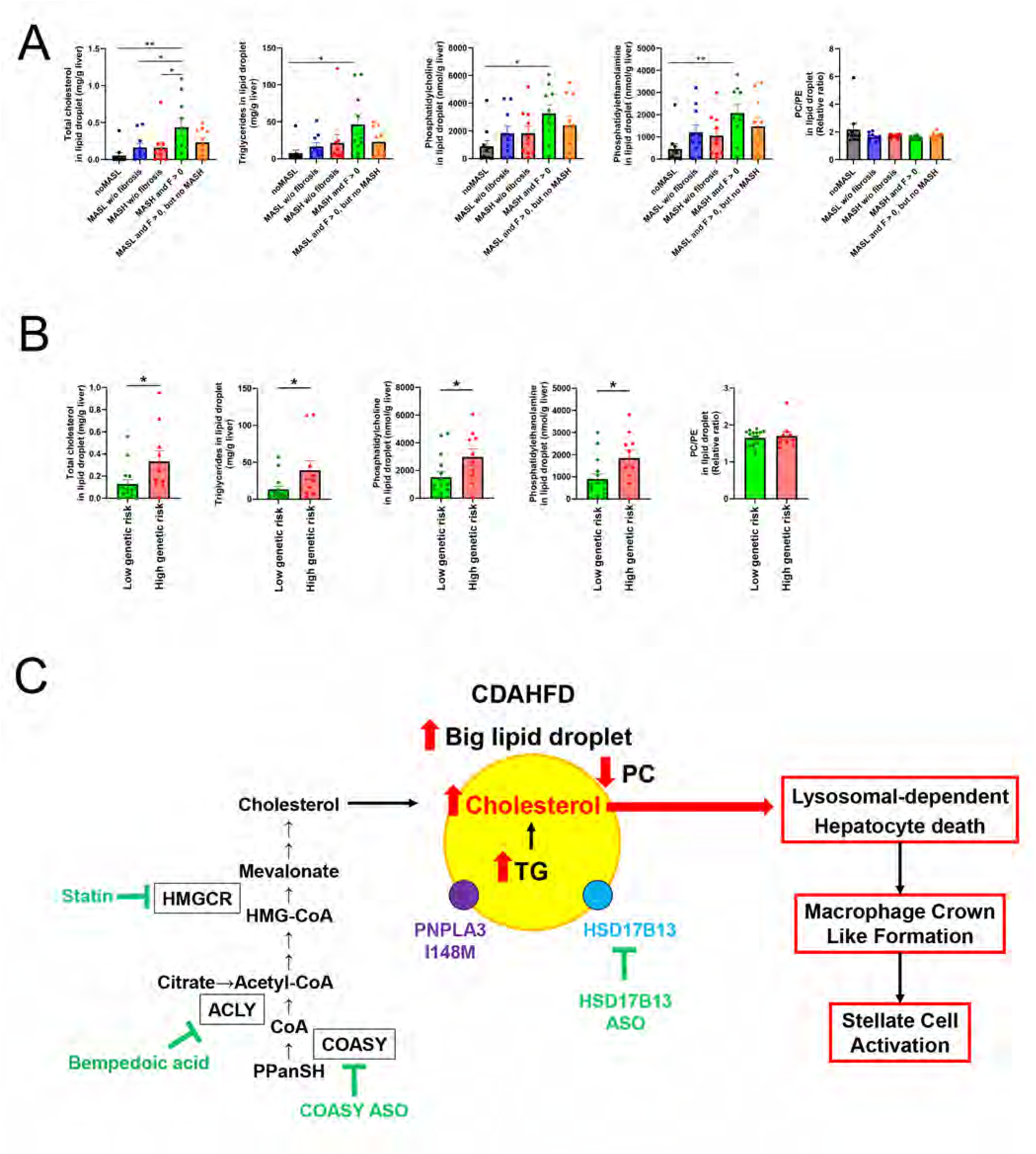
Lipid droplet cholesterol is a key factor in explaining how common genetic variants both cause and protect against metabolic dysfunction-associated steatohepatitis. (A) The metabolite levels in human liver lipid droplets. Total cholesterol, triglycerides, phosphatidylcholine (PC) and phosphatidylethanolamine (PE) in liver lipid droplets were higheset in human metabolic dysfunction-associated steatohepatitis (MASH) with fibrosis compared to other groups. There were no differences in the ratio of PC to PE. Data are presented as mean ± SEM. Groups were compared by one-way ANOVA followed by Tukey’s multiple comparisons test. (B) Comparison of the metabolite in human liver lipid drroplets levels between PNPLA3 I148M variants without HSD17B13 splice site variants (high genetic risk) and HSD17B13 splice variants (rs72613567) without PNPLA3 I148M variants (low genetic risk). (C) The choline-deficient, L-amino acid-defined high-fat diet (CDAHFD) induces the formation of large lipid droplets, including cholesterol, in the liver. Cholesterol accumulation in lipid droplets leads to lysosome-dependent hepatocyte death due to membrane leakage, which results from a relative phosphatidylcholine deficiency on the lipid droplet surface. This process subsequently triggers macrophage crown-like formation and stellate cell activation. Cholesterol-lowering drugs, such as statins and bempedoic acid, exhibit protective effects in the CDAHFD model by reducing cholesterol levels in lipid droplets. Additionally, we identified COASY as a potential therapeutic target for decreasing cholesterol in lipid droplets. Furthermore, cholesterol accumulation in liver lipid droplets contributes to metabolic dysfunction-associated steatohepatitis (MASH) and is influenced by genetic variants, including the PNPLA3 I148M variant and the HSD17B13 splice site variant.

## Discussion

MASLD is the leading chronic liver disease in the world which in turn can progress to MASH and progressive liver fibrosis. However the underlying factors that promote progression from hepatic steatosis to inflammation and liver fibrosis remain unknown (33). As CDAHFD rapidly induces features of MASH in mice, we felt it was an ideal preclinical model to evaluate lipid mediators of hepatic inflammation and fibrosis. We specifically focused on cellular PC, PE, and both total cholesterol content and lipid droplet cholesterol content, as these lipid species have been closely tied to the development of MASH and liver fibrosis. By combining the short-term CDAHFD diet with dietary, molecular, and genetic manipulations we observed that liver lipid droplet cholesterol content was consistently associated with the development of MASH and that inhibitors of the cholesterol synthetic pathway led to reductions in lipid droplet cholesterol content and prevention of the fibrotic response in all of our models.

Several pieces of evidence support a pathogenic role for cholesterol accumulation in MASH (34). In a recent report, MC4R knockout mice fed a Western diet demonstrate that cholesterol accumulation in macrophages plays a key role in the pathological progression of MASH (35). Furthermore, these investigators reported that MASH can be improved by promoting the excretion of macrophage cholesterol using β-cyclodextrin polyrotaxane. In our study, we focused on evaluation of total and free cholesterol content in liver lipid droplets. Total cholesterol consists of cholesterol esters and free cholesterol. Free cholesterol is highly toxic to multiple cellular processes and organelles (36). We also evaluated Filipin staining, which detects free cholesterol. We found that total cholesterol, as well as free cholesterol, increased in liver lipid droplets, which plays a key role in CDAHFD-induced MASH. Accumulation of free cholesterol can form cholesterol crytals that activate inflammation (34). However, in our study, we did not observe any cholesterol crystals in liver lipid droplets by transmission electron microscopy and Filipin staining showed homogenous positive staining of cholesterol in liver lipid droplets suggesting that cholesterol crystallization in lipid droplets is not required for the induction of liver inflammation and fibrosis in this mouse model of MASH.

While cholesterol excess is expected to exacerbate MASH, how cholesterol excess in the liver exacerbates MASH has not been fully elucidated yet. In our study, bempedoic acid or statin treatment decreased liver lipid droplet cholesterol, MASH-associated macrophage markers and liver fibrosis markers (Figure 3 and S3). MASH-associated macrophages were identified using single-cell RNA sequencing (17). This macrophage subset is strongly associated with liver fibrosis (37). Recently, it has been reported that exposure to lipid droplets, released upon injury of steatotic hepatocytes, triggers MASH-associated macrophage induction followed by liver injury (38). In our study, CDAHFD induced large lipid droplets, including cholesterol and crown-like structure, within one week (Figure 1). The crown like structures consist of macrophages surrounding or engulfing dead or dying hepatocytes in MASH; these hepatocytes contain large lipid droplets (11). MASH-associated macrophages are known to be primarily localized to crown-like structures (38). These display a profibrotic phenotype (39). Activated fibroblasts and collagen deposition are observed around crown-like structures, and the number of crown-like structures positively correlates with the fibrosis área (40). These findings suggest that liver lipid droplet cholesterol induces a MASH-associated macrophages/crown-like structure followed by fibrosis.

Furthermore, Coasy knockdown decreased liver lipid droplet cholesterol, MASH-associated macrophage markers and liver fibrosis markers similar to statin or bempedoic acid treatment (Figure 4). COASY converts 4’-Phosphopantetheine (PPanSH) into Coenzyme A (CoA) by adenylation and phosphorylation. COASY is a bifunctional enzyme, the 4th and 5th step of the CoA biosynthetic pathway from Pantothenate (Pan) (41). Whole body loss of function of *COASY* is responsible for an autosomal recessive neurodegenerative disorder but the specific effect of COASY in the liver remains to be clarified (42). Taken together our findings (Figures 4 and 5) indicate that the liver COASY/CoA/acetyl-CoA axis represents a potential therapeutic target to decrease hepatic cholesterol synthesis, leading to MASH and that by targeting COASY knockdown in the liver might avoid the CNS toxicities.

Advances in understanding the genetic underpinnings of MASH have revealed important insights into MASH pathogenesis (27). Protective variants in *HSD17B13* have been identified. PNPLA3 I148M is a risk variant. Accordingly, gene silencing strategies targeting *HSD17B13* and *PNPLA3* are under early phase human clinical trial. Our findings support that the HSD17B13 knockdown strategy could be progressed to human clinical trials through reductions in lipid droplet cholesterol content. Conversely the PNPLA3 I148M accounts for the largest fraction of MASLD heritability (43). PNPLA3 I148M protein was shown to have a loss-of-function by reducing VLDL secretion from hepatocytes and a gain-of-function of accumulation of PNPLA3 on lipid droplets surface leading to suppressed lipid droplet lipolysis (32, 44–47). Our findings indicate that cholesterol in liver lipid droplets plays a key role in the pathogenesis of MASH due to the PNPLA3 I148M variant.

It is reported that phosphorylation of HSD17B13 at serine 33 leads to inhibition of the interaction between CGI-58 and ATGL, followed by reducing intrahepatic lipolysis (48). PNPLA3 148M was shown to compete with ATGL for the interaction with CGI-58, reducing intrahepatic lipolysis (32). Based on these findings, the mechanism of HSD17B13 variants and PNPLA3 I148M are expected to regulate liver lipid droplet lipolysis via ATGL/CGI-58, leading to the alteration of cholesterol in liver lipid droplets.

Furthermore, we confirmed in humans that cholesterol in liver lipid droplets was higher in individuals with MASH and liver fibrosis than in individuals with MASL, MASL without fibrosis, and MASH without fibrosis using liver biopsy samples. Furthermore, consistent with our mouse HSD17B13 gene knockdown and PNPLA3 I148M gene knock-in mouse studies we found that total cholesterol and triglyceride content in liver lipid droplets were higher in PNPLA3 I148M variants without HSD17B13 splice site variants (high genetic risk group) than in HSD17B13 splice site variants without PNPLA3 I148M (low genetic risk group).

Choline deficiency and reduced hepatic PC content have been shown to be associated with liver disease in a variety of conditions (49). Increased liver PC content has been shown to be protective in MASH in carriers of variants HSD17B13 rs72613567 and MARC1 A165T (50, 51) while carriers of the harmful variants PNPLA3 I148M and TM6SF2 E167K demonstrate decreased liver PC content (52, 53). Taken together these results suggest that liver PC content might play a role in the progression of MASH, however the mechanism by which liver PC content alters the pathogenesis of MASH is unknown and prior studies have not evaluated PC content in specific hepatocelular compartments. Here, we focused on PC and PE, specifically in liver lipid droplets, because the lipid droplet membrane consists of a monolayer of PC and PE, which plays a vital role in lipid droplet dynamics (54). PC is cylindrically shaped and would be predicted to provide good coverage of the lipid droplet surface. In contrast, PE is cone-shaped and would be expected to provide less coverage of the lipid droplet surface. We therefore hypothesized that PC deficiency might lead to packing defects on the lipid droplet monolayer (Figure S7) which would then be filled by triglycerides and cholesterol thus exposing these lipid droplet metabolites, which are normally sequestered inside the lipid droplet, to other intracelular organelles such as lysosomes (55).

Free cholesterol accumulation in a gap of phospholipids on the lipid droplet surface is expected to lead to cholesterol crystal formation linked to MASH pathogenesis (36). We confirmed that CDAHFD decreased the ratio of PC to PE in liver lipid droplets and accumulated free cholesterol in liver lipid droplets without cholesterol crystals using Filipin staining and transmission electron microscopy (Figure 2). Based on these findings, free cholesterol accumulation in liver lipid droplets under PC deficiency may lead to free cholesterol leaking from lipid droplets’ surface to other organelles, especially lysosomes, without crystal formation. Transmission electron microscopy revealed lipid droplets in lysosomes and lysosomal membrane disruption (Figure 2). Consistently, cathepsin D immunostaining showed CDAHFD-induced leakage of cathepsin D into the cytoplasm. TUNEL staining revealed apoptotic hepatocytes in CDAHFD-fed mice liver. In general, free cholesterol is highly insoluble, cannot diffuse within the aqueous environment of the lysosomal lumen, and impairs lysosomal function (56). Abnormal cholesterol accumulation in lysosomes induces lysosomal membrane permeabilization, leading to lysosome-dependent cell death (57). It is possible that cholesterol accumulation in liver lipid droplets may cause lysosomal damage followed by lysosome-dependent hepatocyte death linked to CDAHFD-induced MASH pathogenesis.

*Lgals3* mRNA expression was strongly induced by CDAHFD and suppressed by bempedoic acid, statin treatment, *Coasy* ASO or *Hsd17B13* ASO in our study. Cholesterol dietary supplementation or PNPLA3 I148M knock-in increased *Lgals3* mRNA expression. Galectin-3 (gene name Lgals3) is a fibrogenic factor known to be secreted from activated macrophage, leading to hepatic stellate cell activation (20, 58). Galectin-3 inhibitor is now under clinical trial for treating MASH (59). Interestingly, the choline-deficient amino acid-defined diet simultaneously increases *Lgals3* mRNA expression and MASH-associated macrophage markers in mouse livers (60). In addition, *in vitro* analysis using macrophages or smooth muscle cells demonstrates that cholesterol supplementation induces Galectin-3 expression (35, 61). CDAHFD treatment induces liver lipid droplets levels of cholesterol, triggering the development of MASH-associated macrophages, which possibly secret Galectin-3 leading to activation of hepatic stellate cells.

## Conclusion

Taken together, these data identify cholesterol accumulation in liver lipid droplets as a key mediator of liver inflammation and fibrosis in MASH (Figure 7C). The buildup of cholesterol in liver lipid droplets leads to macrophage crown-like formation and stellate cell activation. This mechanism is broadly linked to human MASH pathogenesis and is associated with genetic variants, including PNPLA3 and HSD17B13. Knockdown of COASY and HSD17B13 expression, as well as bempedoic acid treatment, represent novel therapeutic approaches to reducing cholesterol content in liver lipid droplets and preventing the progression of MASLD to MASH and liver fibrosis.

## Materials and Methods

### KEY RESOURCES TABLE

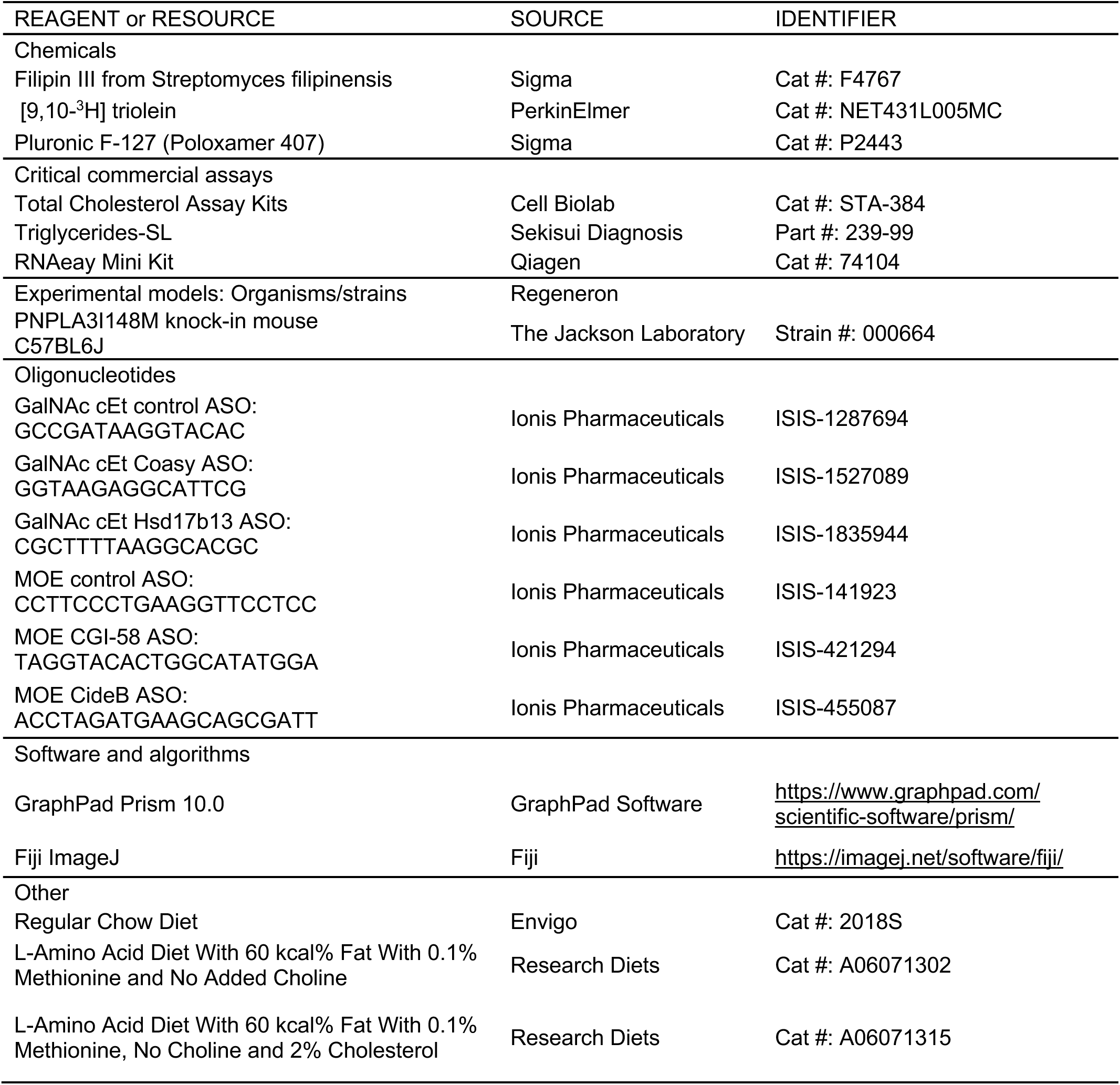

### Animals

The Yale University Institutional Animal Care and Use Committee approved all mouse studies. Male C57BL/6J mice (Jackson Laboratories, Bar Harbor, ME, USA) were group housed at the animal care facility at Yale University Animal Research Center and maintained under controlled temperature and lighting (12:12 h light/dark cycle, lights on at 7 a.m.) with ad lib access to water and food. Mice received weekly intraperitoneal injections of *N*-acetylgalactosamine-modified antisense oligonucleotides (GalNAc-ASO) against *Coasy*, *Hsd17b13* or nontargeting control ASO at a dose of 5 mg/kg per week for three weeks or 2′-O-methoxyethylribose modified antisense oligonucleotides (MOE-ASO) against CideB, CGI-58 or nontargetting control ASO at a dose of 50 mg/kg per week for three weeks. In C57BL/6J mice, a choline-deficient, amino acid-defined high-fat diet (CDAHFD) (60% fat, 0.1% methionine, and no choline, A06071302, Research Diets, New Brunswick, NJ, USA) was used to induce MASH for one day to eight days. CDAHFD with 2% cholesterol (L-Amino Acid Diet With 60 kcal% Fat With 0.1% Methionine, No Choline, & 2% Cholesterol A06071315, Research Diets, New Brunswick, NJ, USA) was used in cholesterol additional treatment study. Mice were treated daily with atorvastatin (20 mg/kg) or bempedoic acid (10 mg/kg) mixed in a small amount (100 mg) of peanut butter or with peanut butter alone as control.

### Human Studies

#### Study population

Liver samples were obtained from 50 obese Caucasian individuals undergoing bariatric surgery from the BARIA_DDZ cohort (ClinTrials.gov identifier: NCT01477957), who were recruited between 2015 and 2023. The BARIA_DDZ cohort study is performed according to the Declaration of Helsinki (updated 2013 version). All participants were informed about the procedures and risks and provided their written consent to the protocol, approved by the ethics boards of Heinrich-Heine-University Düsseldorf and of the Medical Association North Rhine, both Germany (no. 2022-2021_1-andere Forschung erstvotierend / no. 2017222).

## METHOD DETAILS

### Liver lipid droplet fractionation

The subcellular fractionation protocol has been previously described (62). In brief, for the lipid droplet fractionation, 20mg of fresh liver tissue was homogenized in 700ul of homogenization buffer A (20 mM Tris-HCl [pH 7.4], 1mM EDTA, 0.25 mM EGTA, 250 mM sucrose), then 300 μl of 3% sucrose was layered on top of the homogenate. Samples are centrifuged 100k × g for 1hr at 4°C. The lipid cake was completely collected with a needle.

### Lipid metabolites of liver lipid droplets

Tissue lipid metabolites analyses were performed as previously described (6). For measurements of total cholesterol, triglycerides, phosphatidylcholine, and phosphatidylethanolamine, liver lipid droplet extracts from approximately 20 mg of liver tissue were homogenized in 4 mL chloroform/methanol (1:1 vol/vol) with 4 nmoL C17-phosphatidylcholine and C17-phosphatidylethanolamine as internal standard. After phase separation with 1mL dH2O, samples were homogenized and centrifuged, and the methanol/water phase was collected. This supernatant was applied to total cholesterol, triglycerides, phosphatidylcholine, and phosphatidylethanolamine measurements.

To measure total cholesterol, 1 mL supernatant is dried down by nitrogen gas and then diluted by 200 μL 1X Assay Diluent buffer (Total Cholesterol Assay Kits, Cell biolab), including 0.25% NP-40. The supernatant is applied to Total Cholesterol Assay (Total Cholesterol Assay Kits, Cell biolab).

For measurement of triglycerides, 22.5 μL supernatant is dried down by nitrogen gas and then applied to Sekisui’s Triglyceride-SL Reagent Assay (Sekisui Diagnostics, USA).

For measurements of phosphatidylcholine and phosphatidylethanolamine, the supernatant was applied to conditioned Waters Oasis MAX extraction cartridges (Waters Corporation, USA). After a washing step, phosphatidylcholine and phosphatidylethanolamine species were eluted using methanol.

### Liver triglycerides and fatty acids measurement

Approximately 50 mg of liver tissue was homogenized in chloroform/methanol (2:1 vol/vol). Samples were mixed with sulfuric acid (1mol/L) and centrifuged for phase separation. The organic phase was measured using Sekisui’s Triglyceride-SL Reagent (Sekisui Diagnostics, USA). GC/MS determined hepatic fatty acids content.

### Plasma biochemical measurement

Plasma biochemical measurement analyses were performed as previously described (6). Plasma triglyceride was measured using Sekisui’s Triglyceride-SL Reagent (Sekisui Diagnostics, USA). Plasma ALT, AST, and total cholesterol levels were determined by using a COBAS Mira Plus analyzer (Roche, Switzerland).

### Quantitative reverse transcription PCR (RT-qPCR) analysis

RT‒qPCR experiments were performed as previously described (63). The gene-specific mRNA expression values were determined and normalized to those of Arbp as an internal control. Briefly, total RNA (1 μg) was extracted using an RNeasy kit (Qiagen, Valencia, CA, USA) and reverse-transcribed using QuantiTect® Reverse Transcription Kit (Qiagen, Valencia, CA, USA). The cDNA products were subjected to RT-PCR using a Step One Plus Real-Time PCR system (Applied Biosystems, USA). All primer information is provided in Table S3.

### Analysis of data from Gene Expression Omnibus (GEO) database

Hepatic gene expression data in mice liver was collected using GEO RNA-seq Experiments Interactive Navigator (GREIN). Identification number of the dataset is GSE120977 (64).

### Histological analysis

Yale Pathology Tissue Services performed hematoxylin, eosin (HE), and CD68 immunostaining. The number of crown-like structures of CD68 immunostaining were counted using the NIH’s ImageJ. Frozen liver sections were stained with Bodipy and Filipin, identifying lipid droplets and free cholesterol. The slides were subjected to a fluorescence microscope with Bodipy (Excitation 488 nm and Emission > 505 nm) and Filipin (Excitation 735 nm – Two-photon and Emission 420-480 nm).

### Transmission electron microscopy

Transmission electron microscopic examination was performed as previously described (65). Briefly, mouse liver was collected and fixed in 2.5% glutaraldehyde in 0.1 M cacodylate buffer at 4°C overnight. Samples were postfixed in 1% osmium tetroxide in the same buffer for 1 h, stained, dehydrated, and embedded in epoxy resin. Ultrathin sections were stained and examined in a Tecnai 12 BioTWIN electron microscope.

### Liver triglycerides production test

Liver triglycerides production test was performed as previously described (66). Briefly, blood samples were collected after overnight fasting to determine basal plasma triglyceride levels. After basal blood collection, regular chow-fed mice and CDAHFD-fed mice were injected intraperitoneally with poloxamer 407 (1 g/kg of body weight; Sigma-Aldrich, USA) to inhibit LPL activity. Blood samples were collected 1, 2, 3, and 4 h after the injection. The VLDL-triglycerides production rate was calculated by the increase in plasma TG level from baseline.

### Lipid tolerance test

Lipid tolerance test was performed as previously described(67–69). Briefly, RC-fed or CDAHFD-fed mice received a gavage of 10 μL/g corn oil conjugated with 10 μCi of [9,10-^3^H] triolein (PerkinElmer). RC-fed mice were simultaneously gavaged with choline chloride (25 mg/mL in corn oil). CDAHFD-fed mice were not gavaged with choline chloride. Blood was collected by tail vein massage at 0, 1, 2, 3, and 4 h. ^3^H radioactivity in liver tissues was measured by scintillation counting.

### Non-target proteome analysis

Frozen mice liver tissues were redissolved in 100 mM Tris pH 8.0, 4% SDS, 20mM NaCl and 10% acetonitrile using a sonicator (BIORUPTOR II, CosmoBio, Tokyo, Japan). The protein content was measured using a BCA Protein Assay Kit (Thermo Fisher Scientific, Waltham, MA, USA) and adjusted to 150 ng/µl with 100 mM Tris pH 8.0, 4% SDS, 20mM NaCl and 10% acetonitrile. Then 200 µL of protein extract was subjected to clean up and digestion with SP3-LASPmethod (PMID: 38447790) with minor modifications. Briefly, two types of Sera-Mag SpeedBead carboxylate-modified magnetic particles (hydrophilic particles, cat# 45152105050250; hydrophobic particles, cat# 65152105050250; Cytiva, Marlborough, MA, USA) were used. These beads were combined at a 1:1 (v/v) ratio, washed twice with distilled water, and reconstituted in distilled water at a concentration of 10 µg solids/µL. 20 µL of reconstituted beads was then added to the protein extract followed by 1-propanol to bring the final concentration to 75%, with mixing for 20 min. Beads were collected and washed twice with 80% 1-propanol and once with ethanol. Subsequently, the beads were resuspended in 80 µL of 50 mM Tris-HCl (pH 8.0) and 10 mM CaCl_2_ containing 0.02% lauryl maltose neopentyl glycol, and 2 µL of 500 ng/µL Trypsin Platinum (Promega, Madison, WI, USA). Thereafter, to digest the protein, the sample was gently mixed at 37 °C for 14 h. The digested sample was reduced and alkylated with the addition of 8 µL of 110 mM tris(2-carboxyethyl)phosphine and 440 mM 2-chloroacetamide at 80 °C for 15 min, and then acidified with 16 µL of 5% trifluoroacetic acid (TFA). The sample was desalted using a GL-Tip SDB (GL Sciences, Tokyo, Japan), which was washed with 25 µL of 80% acetonitrile (ACN) in 0.1% TFA, followed by equilibration with 50 µL of 3% ACN in 0.1% TFA. The sample was loaded onto the tip, washed with 80 µL 3% ACN in 0.1% TFA, and eluted with 50 µL of 36% ACN in 0.1% TFA. The eluate was dried in a centrifugal evaporator (miVac Duo Concentrator; Genevac, Ipswich, UK). The dried sample was then redissolved in 0.02% decyl maltose neopentyl glycol (DMNG) containing 0.1% TFA. The peptide concentration in the digest sample was measured using a Pierce Quantitative Fluorescent Peptide Assay kit (Thermo Fisher Scientific), according to the manufacturer’s instructions, and adjusted to 200 ng/µL with 0.02% DMNG containing 0.1% TFA.

The peptides were directly injected onto an Global Peptide Separation column (C18, 75 μm ID, 30 cm length, 1.7 µm beads; CoAnn Technologies, Richland, WA, USA) at 50°C and then separated with a 100-min gradient (A = 0.1% formic acid in water, B = 0.1% formic acid in 80% ACN) consisting of 0 min 7% B, 88 min 37% B, 94 min760% B, 100 min 70% B at a flow rate of 150 nL/min using an UltiMate 3000 RSLCnano LC system (Thermo Fisher Scientific). The peptides eluted from the column were analyzed using an Orbitrap Exploris 480 (Thermo Fisher Scientific) with an InSpIon system (AMR, Tokyo, Japan)^86^ for DIA-MS. MS1 spectra were collected in the range of m/z 495–745 at a 60,000 resolution to set an AGC targets of 3 × 10^6^ and a maximum injection time of “Auto”. MS2 spectra were collected at m/z 200–1,800 at a 45,000 resolution to set an AGC targets of 3 × 10^6^, a maximum injection time of “Auto”, and a normalized collision energy of 26%. The isolation width for MS2 was set to 4 Th, window patterns at m/z 500–740 were used for window placements optimized via Scaffold DIA v3.0.1 (Proteome Software., Portland, OR, USA). Six biological replicates were prepared for each of the 16 groups, including one control group, for a total of 96 samples analyzed by LC-MS/MS.

The DIA-MS data were queried against the in *silico* mouse spectral library, using DIA-NN v. 1.8.1(70). Initially, the spectral library was generated from the mouse UniProt protein sequence database (Proteome ID UP000000589, 21949 entries, download on March 7, 2023) using DIA-NN. The parameters for spectral library generation included trypsin as the digestion enzyme, allowance for 1 missed cleavage, a peptide length range of 7–45, precursor charge ranging from 2 to 4, precursor *m*/*z* ranging from 490 to 750, and fragment ion *m*/*z* ranging from 200 to 1800. Additionally, “FASTA digest for library-free search/library generation”, “deep learning-based spectra, RTs, and IM prediction”, “n-term M excision”, and “C carbamidomethylation” were enabled. For the DIA-NN search, the following parameters were applied: a mass accuracy of 10 ppm, MS1 accuracy of 10 ppm, protein inference based on genes, utilization of neural network classifiers in single-pass mode, quantification strategy using robust LC (high precision), cross-run normalization set to “off”. Furthermore, “unrelated runs”, “use isotopologues”, “heuristic protein inference”, and “no shared spectra” were enabled, while the use of match-between-run was deactivated. The threshold for protein identification was set at 1% or less for both precursor and protein false-discovery rate. Protein quantification values were aggregated over the quantification values of unique peptides as calculated by DIA-NN. Differential expression proteins (DEPs) were estimated and analyzed using TCC-GUI. Pathway analysis using Metascape was also performed using proteins with significant DEPs.

#### Liver biopsies

An experienced surgeon obtained the liver biopsies 30 minutes after the induction of anesthesia during bariatric surgery according to a standardized procedure(71). The tissue was immediately snap-frozen in liquid nitrogen and stored at −80°C until further analyses.

#### Liver histology

Histological analysis of liver tissue was performed by a blinded pathologist according to the NASH CRN scoring system(72); MASH was diagnosed when the steatosis grade was ≥1 and combined ballooning/lobular inflammation was present.

#### Genotyping

Genomic DNA was extracted from whole blood with the Qiagen Blood and Tissue kit (Qiagen, Hilden, Germany). Genotyping was performed using real-time-PCR-based allelic discrimination with TaqMan® pre-designed SNP genotyping assays (rs1044498, rs3480, rs1801278, rs6834314, rs695366, rs12137855, rs2642438, rs4374383, rs2294918, rs738409, rs4880, rs58542926, rs641738) and chemistry (ThermoFisher, Darmstadt Germany) on a StepOne Plus Real time PCR System (ThermoFisher). For rs72613567 an assay was designed based on sequence ID NC_000004.12 (87310198–87310323).

| Gene Symbol | SNP ID | Allele | Assay ID |
| --- | --- | --- | --- |
| ENPP1 | rs1044498 | A/C | C_1207994_20 |
| FNDC5 | rs3480 | A/G | C_8822841_10 |
| HSD17B13 | rs72613567 | -/A | custom |
| IRS1 | rs1801278 | C/T | C_2384392_20 |
| KLHL8,MIR5705 | rs6834314 | A/G | C_30687619_10 |
| LARGE1 | rs695366 | T/G | C_2479069_20 |
| LYPLAL1,LOC101929713 | rs12137855 | C/T | C_31403184_10 |
| MARC1 | rs2642438 | A/G | C_1235772_10 |
| MERTK | rs4374383 | G/A | C_26040993_10 |
| PNPLA3 | rs2294918 | A/G | C_2520500_10 |
| PNPLA3 | rs738409 | C/G | C_7241_10 |
| SOD2 | rs4880 | A/G | C_8709053_10 |
| TM6SF2 | rs58542926 | C/T | C_89463510_10 |
| TMC4 | rs641738 | C/T | C_8716820_10 |

## QUANTIFICATION AND STATISTICAL ANALYSIS

The results are presented as the mean ± SEM. Two groups comparisons were performed by unpaired one-tailed Student’s t test. Mupltiple comparisons were performed by one-way ANOVA followed by Tukey’s multiple complarison. Significance is indicated by asterisks, as follows:

*P < 0.05, **P < 0.01; **P < 0.001.

## Acknowledgments

The authors thank John Stack, Xiaoxian Ma, Wanling Zhu and the Yale Histology Core Service for their excellent technical assistance. This study was supported by grants from the United States Public Health Service NIH/NIDDK (F31DK126362 [T.E.L.], T32 GM007324 [T.E.L.], P30 DK34989, R01 DK119968 [G.I.S.], R01 DK113984 [G.I.S.], P30 DK045735 [G.I.S.], R01 DK133143 [G.I.S.]. I.S. was supported by the Manpei Suzuki Diabetes Foundation, Mishima Kaiun Memorial Foundation, Kowa Life Science Foundation and the Ministry of Education, Culture, Sports, Science and Technology (Japan) Fund for the Promotion of Joint International Research (Fostering Joint International Research (A); #20KK0373).

