## Supporting Figures S1 to S7 and Tables S1 to S3 for "Liver lipid droplet cholesterol content is a key determinant of metabolic dysfunction-associated steatohepatitis"

**This PDF file includes:**

Supporting text
Figures S1 to S7
Tables S1 to S3

**Fig. S1. (A) Isolation of lipid droplet fraction from liver. (B) Free cholesterol and cholesteryl ester levels in liver lipid droplets by LC-MS/MS analysis**

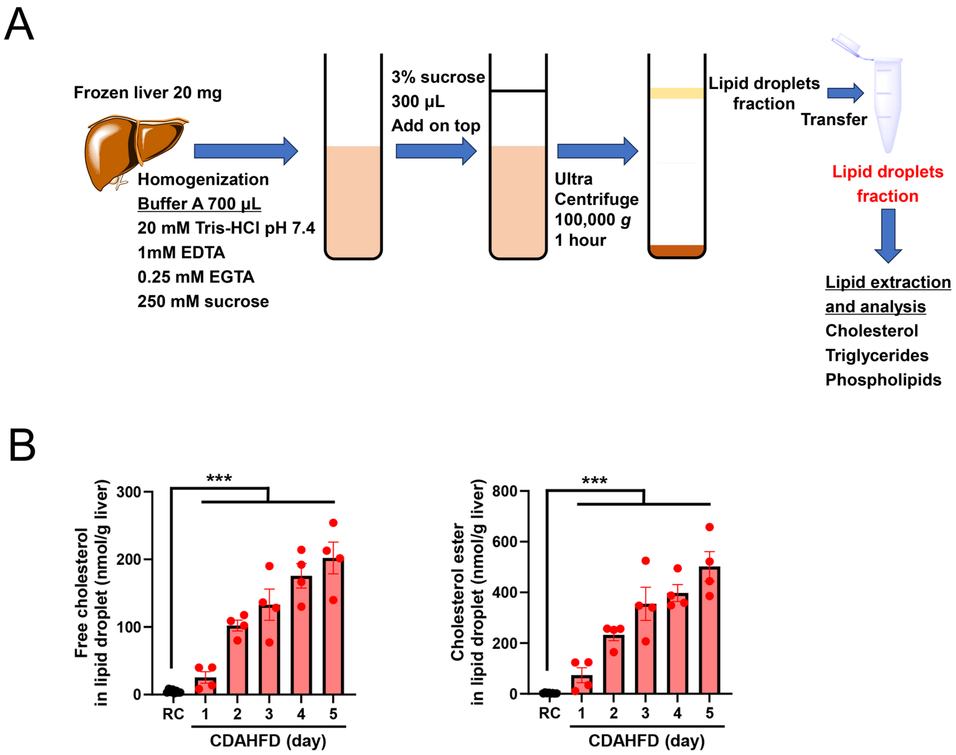

**Fig. S2. (A) Body weight, %fat, %lean mass and plasma glucose levels under six hours fasting of one-week regular chow (RC)-fed mice or choline-deficient, L-amino acid-defined, high-fat diet (CDAHFD)-fed mice related to Figure 2. (B) Hepatic gene expression of CDAHFD 12 weeks fed mice using RNA-sequencing data GSE120977. (Regular chow, n = 5; CDAHFD, n = 5). Transcriptome alterations in 12 weeks of CDAHFD in the GEO database are consistent with the findings of one week model. Data are presented as mean ± SEM. Unpaired one-sided Student's t-test compared groups.**

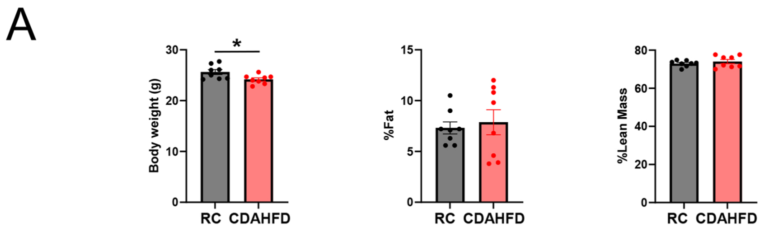


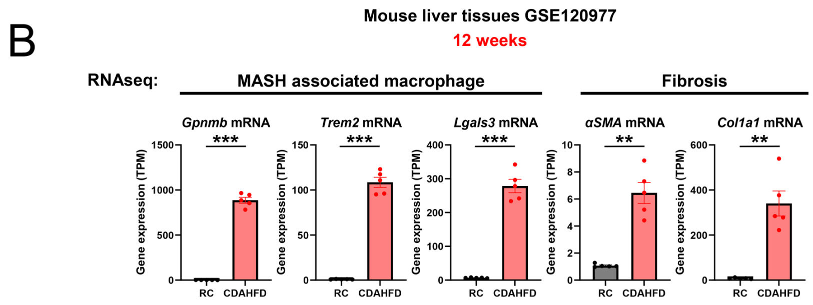

**Fig. S3.** ***Atorvastatin alleviates liver inflammation and fibrosis due to choline-deficient L-amino acid-defined high-fat diet via decreasing cholesterol content in liver lipid droplets.***
**(A) Study design. C57BL/6J mice were divided into a control group and an atorvastatin-treated group. Both groups were fed a choline-deficient, L-amino acid-defined, high-fat diet (CDAHFD) for one week.**
**(B) Plasma ALT and AST in atorvastatin (Statin) treated mice decreased compared to control mice.**
**(C) Liver sections were stained with hematoxylin, eosin (HE), CD68, BODIPY, and Filipin. Statin treatment prevented steatosis. CD68 staining revealed decreased crown-like structures. BODIPY and Filipin staining demonstrated decreased free cholesterol in liver lipid droplet.**
**(D) The number of crown-like structures decreased in statin-treated mice liver.**
**(E) The metabolites levels in liver lipid droplet. Total cholesterol and triglycerides decreased in statin-treated mice. The level of phosphatidylcholine (PC) and phosphatidylethanolamine (PE) did not show a difference. The ratio of PC to PE did not show a difference. The total cholesterol to $PC_{SA}$ ratio decreased in statin-treated mice.**
**(F) Quantitative real-time polymerase chain reaction (RT-qPCR) analysis of liver tissues demonstrated a decreased mRNA expression of MASH-associated macrophage markers (*Gpnmb and Trem2*) and fibrosis markers (*Lgals3, αSMA, and Col1a1*) in statin-treated mice.**
**Data are presented as mean ± SEM. Groups were compared by Unpaired one-sided Student's t-test.**

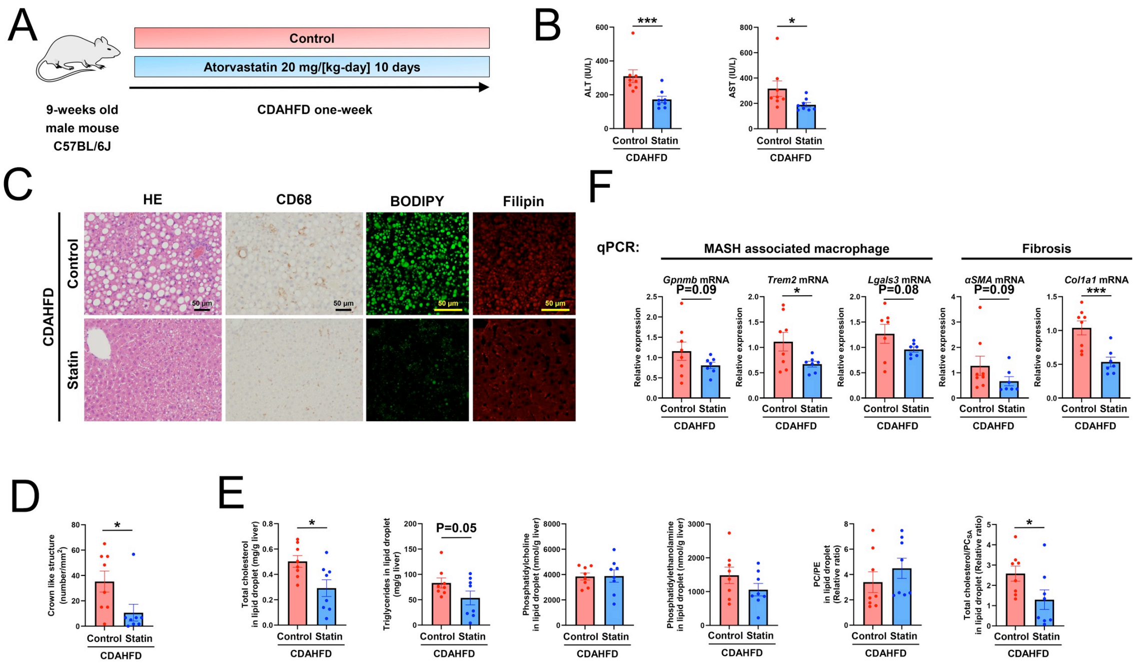

**Fig. S4.** ***Coenzyme A synthase knockdown alleviates liver inflammation and fibrosis due to GAN-diet via decreasing cholesterol in liver lipid droplets.***

**(A) Study design. C57BL/6J mice were divided into GalNAc control ASO-treated mice and GalNAc Coasy ASO-treated mice. Both groups were fed a GAN diet for 40 weeks.**

**(B) Plasma ALT, AST, and liver hydroxyproline content in Coasy ASO-treated mice decreased compared to control ASO-treated mice.**

**(C) The metabolites levels in liver lipidropletset. Total cholesterol and triglycerides decreased in Coasy ASO-treated mice. The level of phosphatidylcholine (PC) and phosphatidylethanolamine (PE) decreased in Coasy ASO-treated mice. The ratio of PC to PE increased.**

**Data are presented as mean ± SEM. Groups were compared by Unpaired one-sided Student's t-test.**

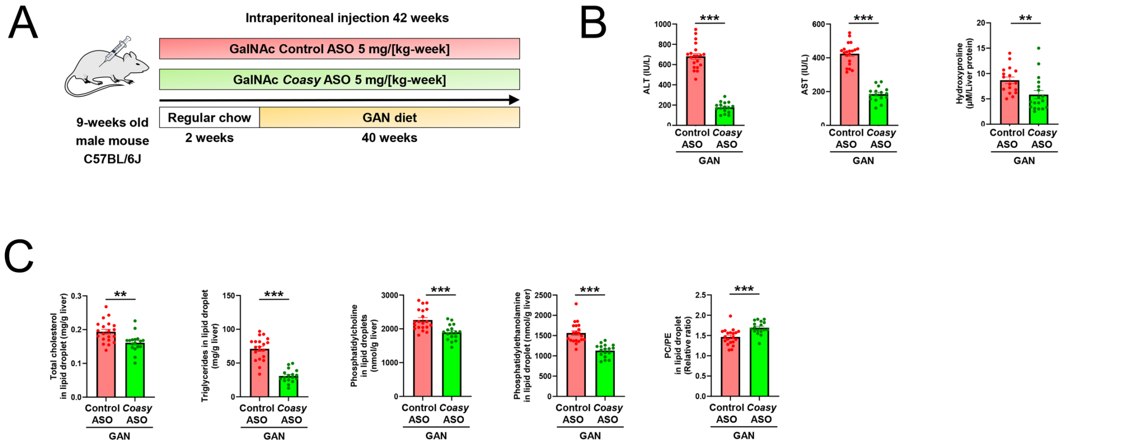

**Fig. S5.** ***Cholesterol supplementation negates the protective effect of Hsd17b13 ASO on a Choline-deficient L-amino acid-defined high-fat diet one-week mice model.***

**(A) Study design. C57BL/6J mice were divided into choline-deficient, L-amino acid-defined, high-fat diet (CDAHFD) treated mice and CDAHFD with 2 % cholesterol treated mice. Both groups were injected with GalNAc *Hsd17b13* ASO.**

**(B) Cholesterol (Chol) dietary supplementation increased plasma ALT and AST levels compared to control (Ctr) mice.**

**(C) Liver sections were stained with hematoxylin, eosin (HE) and CD68. Dietary cholesterol supplementation induced steatosis. CD68 staining revealed increased crown-like structures.**

**(D) The number of crown-like structures increased in dietary cholesterol-supplemented mice liver.**

**(E) The metabolites levels in liver lipid droplet. Total cholesterol and triglycerides increased in cholesterol-overload mice. The level of phosphatidylcholine (PC) and phosphatidylethanolamine increased (PE) in cholesterol-supplemented mice. The ratio of PC to PE remained unchanged.**

**(F) Quantitative real-time polymerase chain reaction (RT-qPCR) analysis of liver tissues showed an elevated mRNA expression of *Gpnmb, Trem2 and Col1a1* in cholesterol-overload mice.**

**Data are presented as mean ± SEM. Groups were compared by Unpaired one-sided Student's t-test.**

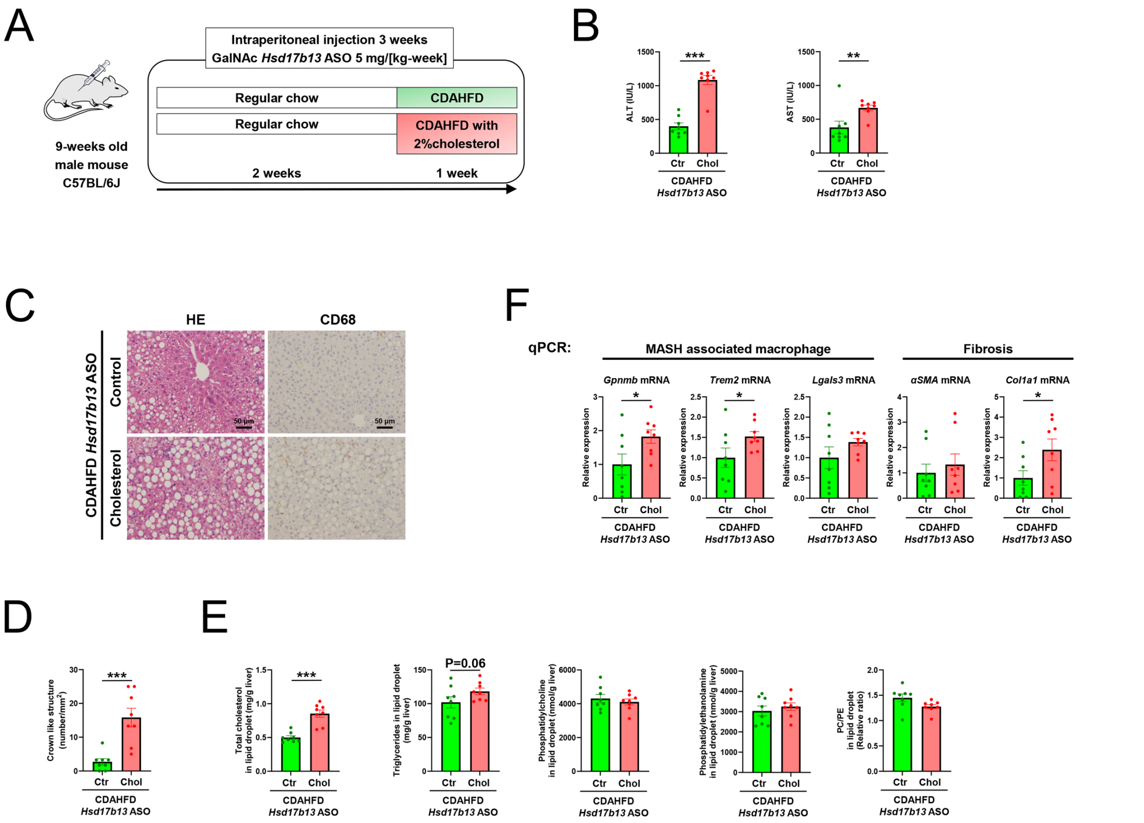

**Fig. S6.** ***Bempedoic acid alleviates liver inflammation and fibrosis of PNPLA3 I148M knockin mice fed with choline-deficient L-amino acid-defined high-fat diet via decreasing cholesterol content in liver lipid droplets.***

**(A) Study design. PNPLA3 I148M knock in mice were divided into a control group and bempedoic acid-treated group. Both groups were fed a CDAHFD for one week.**

**(B) ALT and AST in bempedoic acid (BemA) treated mice decreased compared to control mice.**

**(C) Liver sections were stained with hematoxylin, eosin (HE) and CD68. BemA treatment prevented steatosis. CD68 staining revealed decreased crown-like structures.**

**(D) The number of crown-like structures decreased in BemA-treated mice liver.**

**(E) The metabolites levels in liver lipid droplet. Total cholesterol and triglycerides decreased in BemA-treated mice. The level of phosphatidylcholine (PC) ande phosphatidylethanolamine (PE) did not show a difference. The ratio of PC to PE did not show a difference.**

**(F) Quantitative real-time polymerase chain reaction (RT-qPCR) analysis of liver tissues demonstrated a decreased mRNA expression of MASH-associated macrophage markers (*Gpnmb*, *Trem2*, and *Lgals3*), and fibrosis markers (αSMA and *Col1a1*) in BemA-treated mice.**

**Data are presented as mean ± SEM. Groups were compared by Unpaired one-sided Student's t-test.**

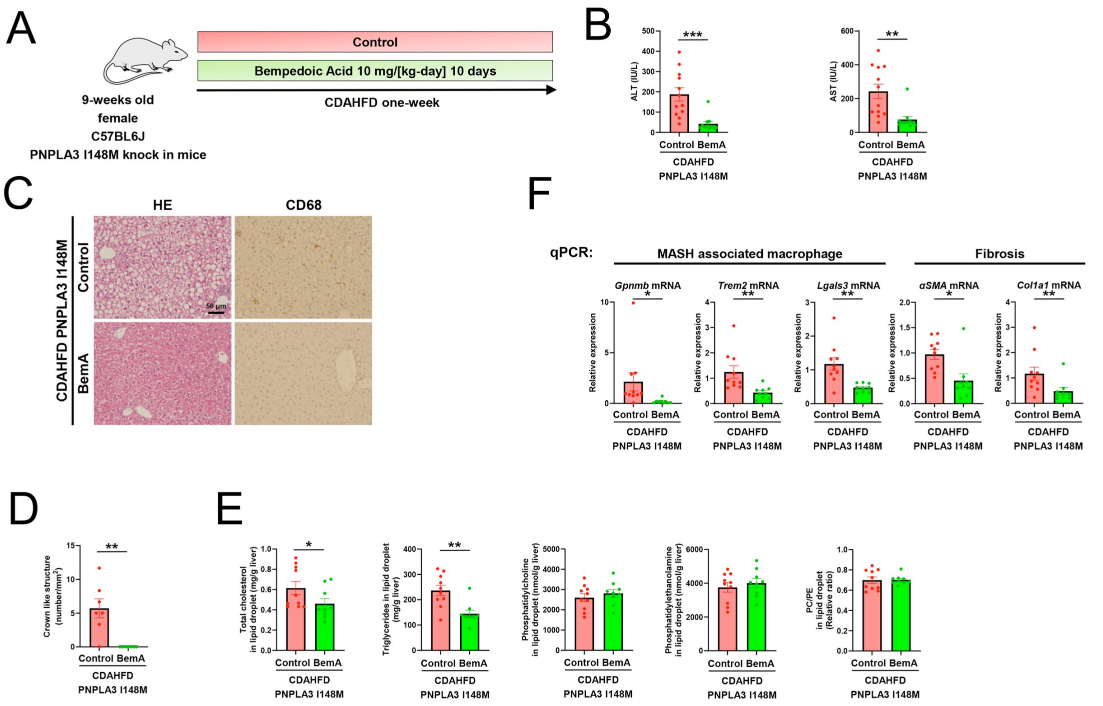

**Fig. S7. (A) Packing defects under phosphatidylcholine deficiency on lipid droplet surface.**

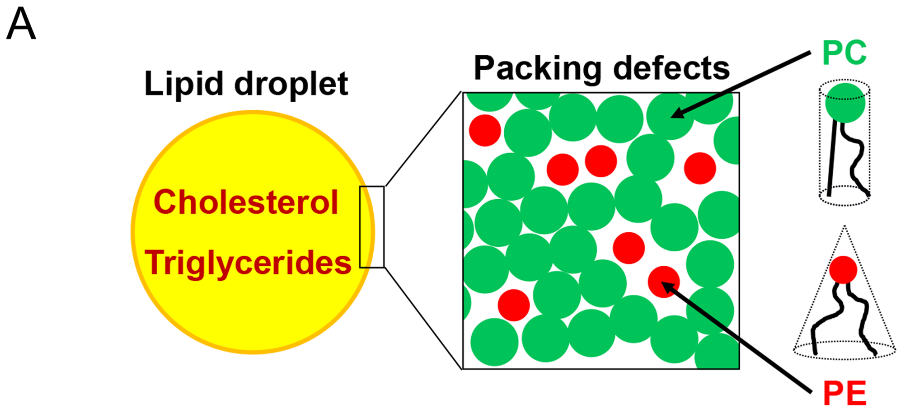

**Table S1.** Human liver samples and genotyping

| | Number | PNPLA3 I148M (C>G)<br>rs738409 | HSD17B13 TA<br>rs72613567 |
|---|---|---|---|
| no MASL | n=10 | I/I=7, I/M=3, M/M=0 | T/T=6, T/TA=3, TA/TA=1 |
| MASL without fibrosis | n=10 | I/I=7, I/M=3, M/M=0 | T/T=6, T/TA=4, TA/TA=0 |
| MASH without fibrosis | n=10 | I/I=5, I/M=3, M/M=2 | T/T=4, T/TA=4, TA/TA=2 |
| MASH and F>0 | n=10 | I/I=6, I/M=3, M/M=1 | T/T=6, T/TA=4, TA/TA=0 |
| MASL and F>0, but no MASH | n=10 | I/I=4, I/M=5, M/M=0, no data=1 | T/T=3, T/TA=6, TA/TA=0, no data=1 |

**Table S2.** Distribution of PNPLA3 rs738409 and HSD17B13 rs72613567 in human liver samples

| | | PNPLA3 I148M rs738409 | |
|---|---|---|---|
| | | II | IM/MM |
| HSD17B13 | TT | 15 | 10 |
| rs72613567 | TTA/TATA | 15 | 9 |

**Table S3.** Primers used for quantitative reverse transcription PCR analysis.

| Gene | | Primers | PCR product size (bp) |
|---|---|---|---|
| *mouse Gpnmb* | Forward | GCTGGTCTTCGGATGAAAATGA | 156 |
| | Reverse | CCACAAAGGTGATATTGGAACCC | |
| *mouse Trem2* | Forward | CTGGAACCGTCACCATCACTC | 183 |
| | Reverse | CGAAACTCGATGACTCCTCGG | |
| *Mouse Lgals3* | Forward | AGACAGCTTTTCGCTTAACGA | 210 |
| | Reverse | GGGTAGGCACTAGGAGGAGC | |
| *Mouse αSMA* | Forward | GAGACTCTCTTCCAGCCATCT | 129 |
| | Reverse | CCCTGACAGGACGTTGTTAGC | |
| *Mouse Col1a1* | Forward | CGATGGATTCCCGTTCGAGT | 197 |
| | Reverse | CGATCTCGTTGGATCCCTGG | |
| *Mouse Coasy* | Forward | ATTTCATCACGCACCTCTACAC | 219 |
| | Reverse | AACTGGTGGCATAACGCTCTA | |
| *Mouse CideB* | Forward | CAATGGCCTGCTAAGGTCAGT | 101 |
| | Reverse | GATCACAGACACGGAAGGGTC | |
| *mouse CGI-58* | Forward | TGGGGTTTTCCTGAGCGAC | 94 |
| | Reverse | GGTTAAAGGGAGTCAATGCTGC | |
| *mouse Hsd17b13* | Forward | ATTCCCCGGAGAAGGAAATCT | 75 |
| | Reverse | CAGCCTGCCTATTCCGTGT | |
| *mouse Arbp* | Forward | GAGGAATCAGATGAGGATATGGGA | 72 |
| | Reverse | AAGCAGGCTGACTTGGTTGC | |
